# Gain of Function p53 mutant R273H confers distinct methylation profiles and consequent partial or full EMT states to colon tumour

**DOI:** 10.1101/2025.07.03.662954

**Authors:** Harsha Rani, Seemadri Subhadarshini, Mohit Kumar Jolly, Vijayalakshmi Mahadevan

**Affiliations:** Institute of Bioinformatics and Applied Biotechnology (IBAB), Bangalore, India; Manipal Academy of Higher Education, Manipal, India; Department of Bioengineering, Indian Institute of Science, Bangalore, India

## Abstract

p53 is the second most frequently mutated gene in colorectal cancer. While different p53 mutations have been correlated with metastasis, the distinct phenotypes exhibited by site-specific mutations of p53 are not well elucidated. Here, we analyse transcriptomic and methylation data from TCGA-COAD cohort to understand the epigenetic impact of three most prevalent hotspot mutations of p53 (R175H, R273H and R282W). We observed that p53 R273H mutation associates with a partial epithelial-mesenchymal transition (pEMT) state and metastatic progression. In vitro ChIP-seq experiments conducted on p53R27H harbouring HT29 cells revealed an enrichment of mutant p53 R273H at pEMT or mesenchymal gene sets. Further, simulations from a gene regulatory network incorporating the interactions of p53R273H with EMT regulators explain how this mutation shapes the phenotypic landscape accessible to cancer cells. Finally, single-cell transcriptomic analysis of colorectal tumours reveals R273H-linked enrichment of partial and mesenchymal EMT phenotypes across tumour subpopulations in CRC. Overall, we identified distinct epigenetic regulation regulating partial EMT and consequent aggressive behaviour triggered by p53R273H. These findings can help devise effective therapeutic strategies for p53 mutant specific colon tumours.

**Graphical Abstract Caption.**
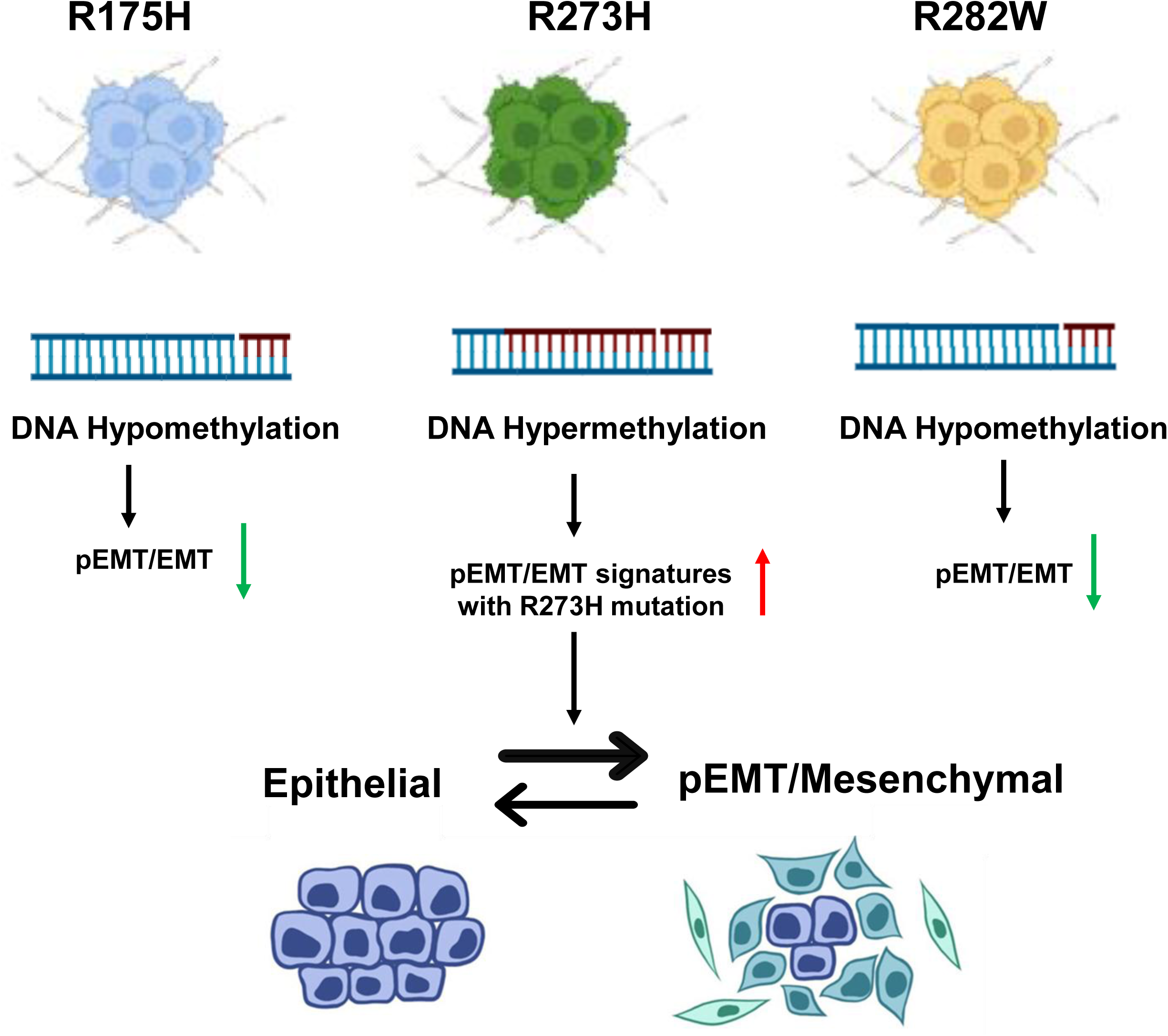
Mutation in Tp53 at R273H associates with DNA hypermethylation and elevated pEMT and metastatic signatures as compared to R175 and R282W mutations in colon tumours suggesting of devising novel therapeutic intervention strategies based on p53 mutation profiles.

## Introduction

The tumor suppressor TP53 is one of the most frequently altered tumour suppressor genes across various cancers. These mutations result in loss of tumour suppressor function of p53 and are known to confer oncogenic “gain-of-function” properties. Recent work in the field has established that 70% of mutations in colorectal cancer is associated with p53 [1,2] and function as driver mutations leading the progression of CRC from late adenoma stage to the carcinoma stage [3, 4]. Mutations in p53 are known to be aggressive and show poor prognosis making mutant p53 a potential and viable target for therapy.

The mutations in p53 encompass truncating and splice sites mutations, in-frame insertions/deletions (indels), frameshift indels and missense mutations. Missense mutations represent the vast majority of mutations occurring in different codons of the TP53 gene (>80%), with the highest frequency in the DNA-binding domain. These mutations lead to deactivation of the tumor suppressor role of p53 and gain of novel oncogenic functions including cell proliferation, antiapoptotic effects, and metastasis formation [5,6] and are therefore defined as gain-of-function (GoF) mutations, a subtype of TP53 mutations, which are clustered as hotspots in the DNA binding domain of Tp53.

International Agency for Research on Cancer (IARC) includes R175, G245, R248, R249, R273, and R282 as the top six frequently mutated sites in Tp53 [7]. Structural studies of the p53-DNA complex have identified that the hotspots mutations at the R248 and R273 amino acid position influence the direct contact with DNA, whereas R249 and R282 lie near the DNA contacting interface [8]. The R175H mutations, one of the most abundant hotspot mutations at Tp53 are located at the zinc-binding site near the DNA binding interface. These hotspot DNA mutants of Tp53 are further classified into contact mutations (R282W, R273H) or conformational mutations (R175H, G245S, and R249S) [9].

Clinical studies, generally considers all *TP53* mutations as collective mutation without any differentiation between the various types of aberration, however, several studies on *in vivo and in vitro* model systems, have shown different phenotypes associated with different p53 mutations [10, 6, 11]. Each mutation differentially affects the survival outcomes of the patients. The phenotype associated with a distinct mutation of Tp53 thus becomes an important deterministic factor for the chemotherapeutic regimen.

A more recent work elucidating the mutant-specific impact of p53R270H and p53R172H on metabolism in pancreatic cancer has revealed unique transcriptomic profiles with p53R270H and p53R172H mutations. p53^R270H^ induces alterations in the expression of genes associated with oxidative stress and reduction in mitochondrial respiration while p53^R172H^ specifically impacts the expression levels of enzymes involved in the urea metabolism [12]. Furthermore *in-vitro* and *in-vivo* studies have shown that p53-R273H exhibits more invasive ability than p53-R175H and p53-R248W and promotes colon cancer stemness expansion [13]. The varied phenotype with diverse p53 mutation spectrum makes it crucial to understand the regulatory mechanisms that drive oncogenesis and progression.

Epigenetic mechanisms including DNA methylation, non-coding RNA and post translational mechanisms of histones play key roles in regulating cancer progression. It is therefore important to understand the epigenetic regulation of mutant and WT p53 signalling to derive newer insights into cancer therapy. Tp53 hotspot mutations R175H, R248Q, R248W, R249S, and R273H transcriptionally upregulate chromatin regulatory genes including methyltransferases MLL1 (KMT2A), MLL2 (KMT2D), and acetyltransferase MOZ (KAT6A) by binding to their promoter regions together with ETS2 [14], suggesting that mutant p53 regulates the different phenotypes by regulating the expression of various epigenetic factors and by co-opting chromatin pathways.

Aberrant DNA methylation of CpG islands has been reported in the earliest detectable lesions in the colonic mucosa. Colorectal tumors with CpG island methylator phenotypes (CIMP) exhibit a high frequency of cancer-specific DNA hyper methylation on a subset of genomic loci and are highly enriched for an activating mutation of BRAF. DNA hypermethylation of some CIMP-associated gene promoters has been detected in early stages of colorectal tumorigenesis. However, only about 11% of Tp53 mutations are associated with CIMP(+ve) phenotype [15]. WT tumor suppressor protein p53 has been shown to cooperate with DNA methylation to maintain silencing of a large portion of the mouse genome[16].

Loss of p53 in colon cancer is associated with an increase in expression of DNMT1 [17]. More recent work using pancreatic cancer as a model system has highlighted that p53 deficiency leads to a reduction in the universal methyl donor (S-adenosyl methionine) SAM, suggesting that loss of p53 fails to maintain epigenetic integrity mainly due to the inability to restore constitutive heterochromatin [18]. A significant correlation has also been observed between the expression of miR-145 and the status of p53 gene in both prostate tissues and 47 cancer cell lines. miR-145 expression was found to be downregulated with varying status of p53 mutation in prostate cancer cell lines. The repression of these loci was regulated by DNA methylation at the CpG sites [19]. Although, direct association of DNA methylation with WTp53 has been investigated, the alterations in DNA methylation level with respect to different hotspot mutations of Tp53 has not been explored.

Mutant p53 is also implied in cancer metastasis [20, 21]. Furthermore, recent studies have demonstrated that that EMT transition is not a binary process, but occurs through distinct cellular states [22, 23], with the hybrid EMT state (pEMT/partial EMT) presenting the highest metastatic potential. This intermediate E/M stages are characterized by different levels of E and M markers, transcriptional, and epigenetic reprogramming which promotes more invasiveness [23]. P-cad and Slug are indicated to play a pivotal role in hybrid EMT and collective cell migration. However, the connection between GoF/LoF mutations in and genes contributing to pEMT states has not been well established.

This work attempts to explore the phenotypes associated with three prevalent hotspot mutations of p53 (R175H, R273H and R282W) and to identify epigenetic signatures driving these distinct phenotypes using transcriptomic profiling and DNA methylation of colon cancer patients from the TCGA-COAD cohort. We have identified distinct transcriptomic profiles regulating each of the p53 mutations considered in the study. Single sample gene set enrichment analysis(ssGSEA) on the transcriptomic signatures with different Tp53 mutation reveals distinct reactome pathways associated with each of the mutation, with R273H to be enriched with YAP/TAZ signalling while R175H with WDR5 histone containing complexes suggesting different regulatory mechanism governing each of these mutations. The observed YAP/TAZ activation in Tp53R273H is indicative of increased cancer stemness and self-renewal contributing towards initiation and progression as compared to R175H and R282W mutations. CMS classification of each of the different p53 mutations reveals distinct enrichment of profiles with R282W and R273H and to have higher CMS4 correlation as compared to R175H. Significant increase in the expression profile of DNMT3A further suggests an epigenetic regulation accompanying the distinct transcriptional signatures with Tp53 mutation. The DNA methylation profiles of the colon cancer patients reported in the cohort further reveals novel methylome profiles with different Tp53 mutation, with R273H mutation accompanying hypermethylated genome which is hitherto unestablished. Hypomethylated genes accompanying R273H mutation were associated with unique upregulated Epithelial to Mesenchymal transcription factors (EMT-TFs) which suggest an epigenetic control governing cancer stemness and metastasis.

ChIP sequencing of p53 mutant cell lines and the Gene Regulatory Network (GRN) establishes that R273H promotes cancer progression through favoring hybrid and metastasis state. These observations are further validated through single cell transcriptome profile in CRC patients. Overall, this work for the first time has identified distinct epigenetic regulation contributing towards pEMT states and metastatic transition upon R273H. These findings reveal novel biomarkers and suggest a novel therapeutic intervention for different tumours with p53 mutational status.

## Results

### p53 mutant colon cancer patients show distinct mutation dependent phenotypes

Mutations on the DNA Binding Domain of p53 have been reported to confer oncogenic and neoplastic functions in various cancer types. These mutations include truncating and splice sites mutations, in-frame insertions/deletions (indels), frameshift indels and missense mutations. Our exploratory analysis of the cBioportal involving 15 different studies from 4233 colorectal patients (including 2936 TCGA colorectal cancer patients) reveals Tp53 mutations (2293 cases) to be the second most mutated gene following APC (Fig S1A). The most prevalent mutation amongst them is missense mutation, followed by truncating mutations (Fig S1B and S1C).

Out of the total 459 colon cancer patients explored from the TCGA-COAD project, 237 patients (51.6%) showed mutations in TP53, with the most prevalent mutation being the missense mutations (68.30%) (Fig S1B, S1C). Among these missense mutations, the conformational mutation R175H and contact mutations R273H and R282W were the most prevalent in the cohort considered (Fig 1A).

**Fig 1:**
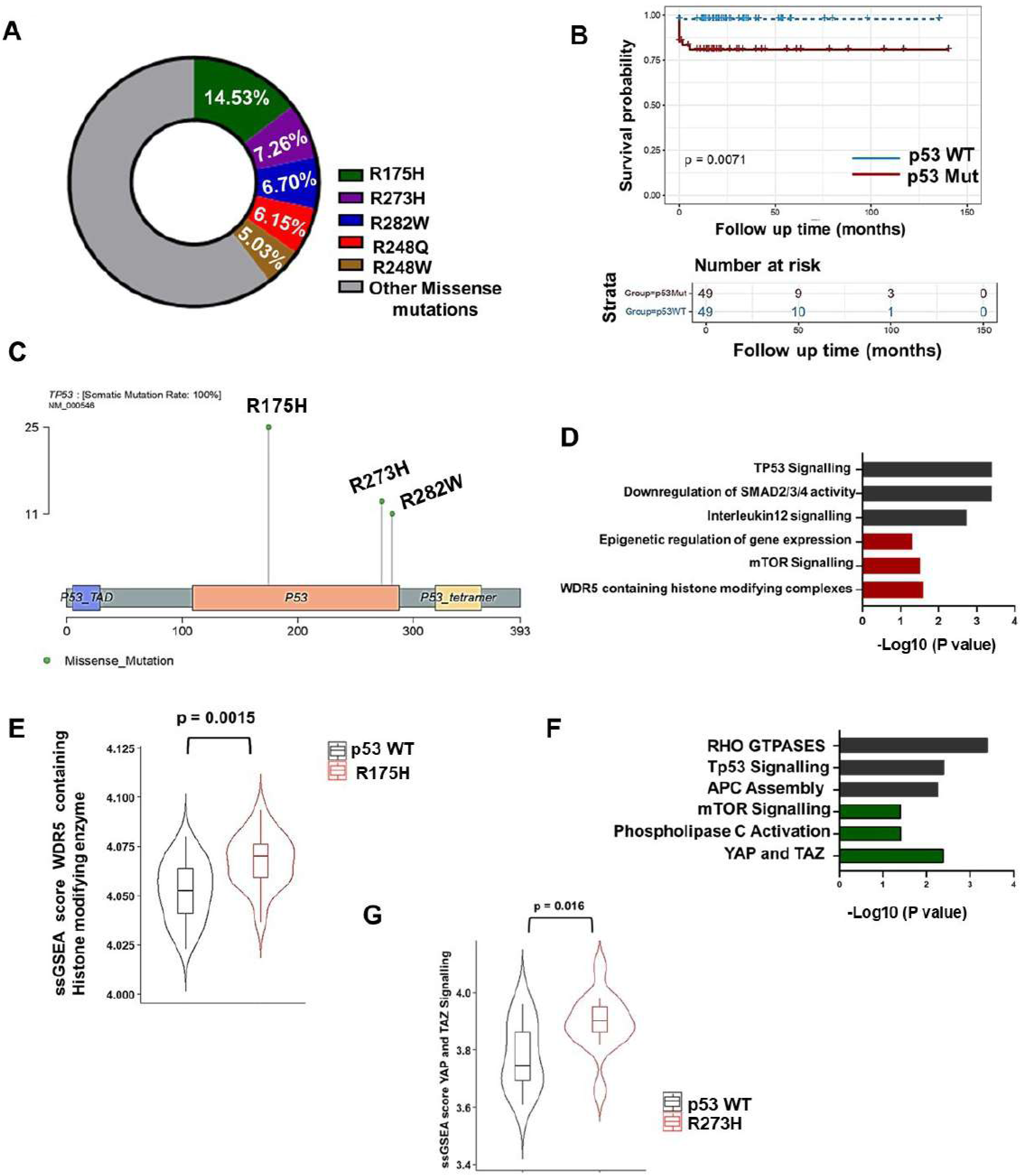
Different p53 missense mutation shows distinct phenotype in colon cancer. A. Distribution of different Tp53 missense mutations in colon tumours as observed in the TCGA-COAD cohort considered for the study. The top three missense mutations in Tp53 include R175H, R273H and R282W. B. Kaplan Meier (KM) plot representing the survival profiles of Tp53WT tumours and top 3 prevalent Tp53 missense mutations (R175H, R273H and R282W). Tumours with Tp53 missense mutations associate with lower survivability as compared to p53 WT tumours [p value <=0.05]. The X axis represents the survival time in months while the Y axis represents the survival probability C. Lollipop plot representing different sites of p53 mutation in p53 missense tumours through COAD cohort. Identification of tumours with p53 R175H tumours (N=25), p53 R273H tumours (N=13) and p53 R282W tumours (N=11). Tumours harboring substitution mutation at Tp53 at the 175^th^ amino acid position where the Arginine is replaced by Histidine, at the 273th amino acid position where the Arginine is replaced by Histidine and at the 282 position where the Arginine is substituted with Tryptophan D. Column chart representing Gene Set Enrichment Analysis (GSEA) of the differentially expressed genes in p53R175H tumours(red) and p53 WT (grey) (through reactome). R175H tumours show positive enrichment of WDR5 containing histone complexes. WDR5 (WD repeat domain 5) functions as an epigenetic reader which recognizes specific histone modifications. E. Box and Violin plots representing distribution of single sample gene set ssGSEA score of WDR5 containing histone modifying enzymes in p53 WT tumours(black) and R175H tumours(red). Tumours with the R175H mutation show higher enrichment score for WDR5 containing histone modifying enzymes F. Column chart representing Gene Set Enrichment Analysis (GSEA) of the differentially expressed genes in R273H tumours (green) and p53 WT (grey) tumours(through reactome pathways). R273H tumours show positive enrichment of YAP/TAZ signalling pathway which promotes epithelial to metastatic transition. G. Box and Violin plot representing distribution of ssGSEA score of YAP/TAZ in p53 WT tumours(black) and R273H tumours(red). Tumours with R273H tumours shows higher enrichment score for YAP/TAZ

We first identified tumours from the TCGA colon cancer cohort, harboring three most prevalent gain-of-function missense mutations (or hotspot mutations) in the DNA-binding domain of p53 at distinct amino acid position (R175H, R273H and R282W), and profiled the transcriptome of these patients (Fig 1C). We similarly identified p53WT colon tumours for which no mutation in Tp53 was found (based on somatic mutation profile) which were used as control data sets for the differential expression analysis from RNA sequencing data.

We examined the influence of these mutations on overall survival of the patients and observed that colon tumours harboring these missense mutations show poor survivability as against p53WT tumours (Fig 1B). We further compared the survival rate of patients with colon tumours harbouring distinct p53 mutations (Fig S1D) and observed that all the three missense mutations are associated with different survival rates.

To understand the diverse phenotypes and transcriptional regulatory functions of each of the p53 mutations, we performed differential gene expression analysis from RNA sequencing of these patients and gene set enrichment analysis (GSEA) and single sample gene set enrichment analysis (ssGSEA) using REACTOME pathways [24]. Interestingly, we observed distinct pathways associated with different p53 mutations with YAP-TAZ signalling pathway to be associated with R273H mutation (Fig 1D-G) and WDR5 epigenetic regulations to be associated with R175H mutation, while p53 related signaling pathways appeared among most of the significantly enhanced pathways in wild-type TP53 cancers involved (Fig S1E).

### p53 mutant R273H regulates YAP dependent metastasis in colon cancer patients

We identified more epigenetically driven reactome pathways (Fig1 and FigS1E) to be associated with R175H mutation suggesting a key distinct role of various WDR5 mediated epigenetic regulators in silencing/activation of tumour suppressors and oncogenes. WT p53 functions as a negative regulator for Yes Associated Protein (YAP). *Invivo and invitro* studies have shown that mutant p53 interacts with YAP1 complex and promotes their pro-proliferative transcriptional activity via activation of NF-Y [25]. More recent study using CRC cells as a model system has shown that YAP1 can inhibit autophagy in human CRC cells by transcriptionally upregulating *Bcl-2*, and thus promote CRC progression [26]. Gene set Enrichment analysis and ssGSEA for genes upregulated with R273H mutation patients suggest YAP1- and WWTR1 (TAZ)-stimulated gene expression. These observations suggest that distinct transcriptional programs are associated with R273H mutation (Fig1G and FigS1F) and a significant enrichment of R273H with hippo signalling pathway, cancer stemness and self-renewal properties as compared to R175H and R282W.

### Colon tumours harbouring R273H mutation alter the transcription landscape of EMT and pEMT

Malignant tumours are often characterized by partial EMT (pEMT) states which involve upregulation of certain mesenchymal genes and moderation of epithelial programs (Pastushenko & Blanpain, 2019, Sinha et al., 2020).Although Gain-of-function (GoF) Tp53 mutation together with loss of wild-type *Tp53* (by loss of heterozygosity (LoH) was found to be a key genetic event for activin A–induced partial EMT in colon cancer [27], the comparative expression profiles of hallmark EMT genes and partial EMT genes with different p53 mutation status have not been studied yet. We therefore compared the expressions of hallmark EMT genes and partial EMT (pEMT) genes identified in the previous study [28] with different Gain of Function (GoF) mutations. We identified R273H mutation to be associated with significantly higher expression of hallmark EMT gene signatures and pEMT genes as compared to R175H and R282W mutations (Fig 2A, B). This observation suggests that R273H preferentially alters the EMT and pEMT states by reprogramming the transcriptional landscape and elevating the expression of genes involved in pEMT and mesenchymal states.

**Fig 2:**
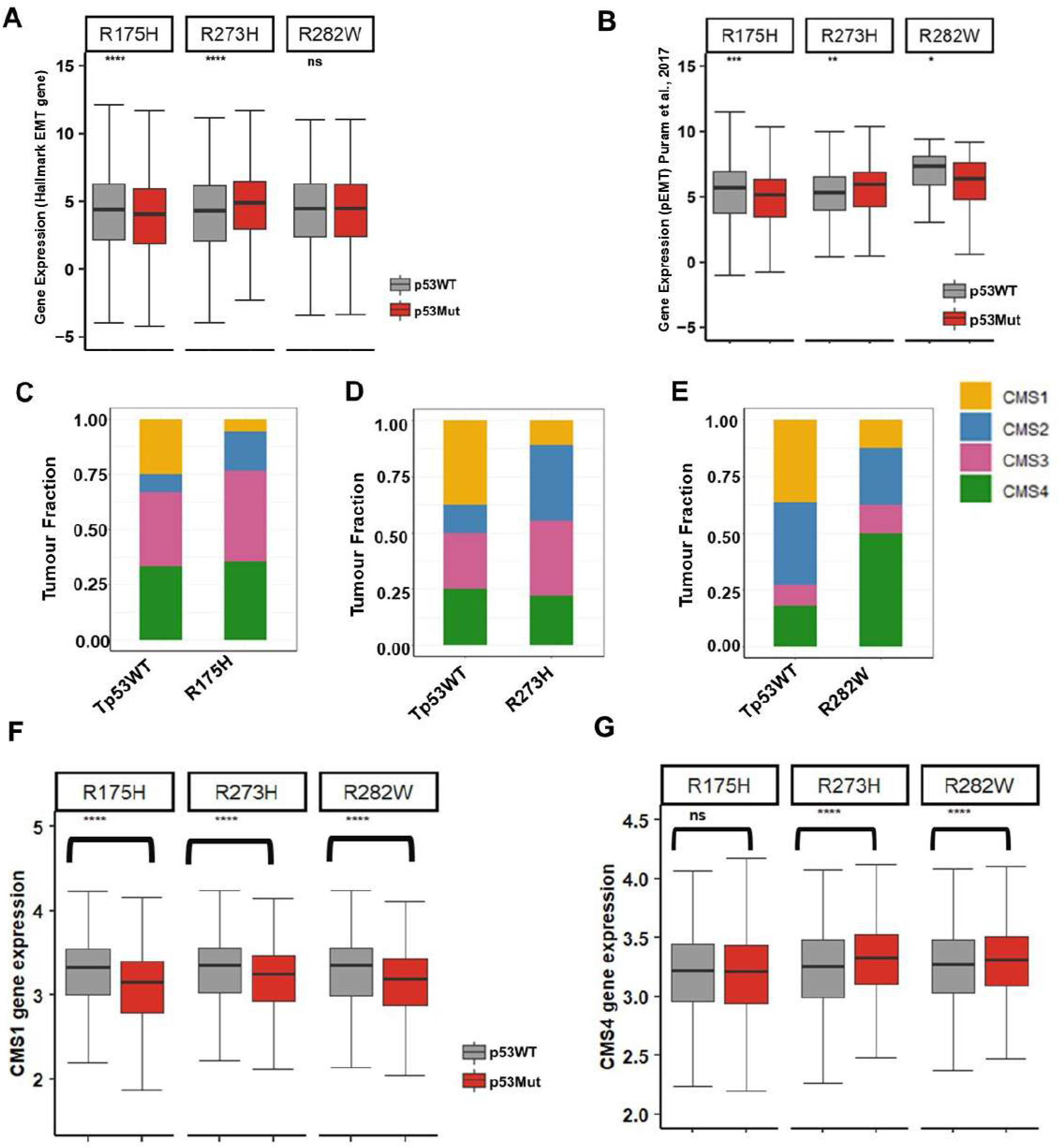
Different p53 missense mutations show varied proportion of CMS subtypes and their associated gene expression. A. Box plot representing the distribution of Hallmark EMT genes in p53WT (grey) and p53mut (red). R273H mutation in Tp53 associates with a significantly higher expression of EMT gene signatures. B. Box plot representing the distribution of expression of partial EMT genes in p53WT (grey) and p53mut(red). R273H mutation in Tp53 associates with a significantly higher expression of Mesenchymal genes C. Stacked bar plot representing CMS classification in WT p53 and p53 R175H COAD cohort (based on CMS classifier). Patients with R175H show higher proportion of CMS3 subtype (pink). The Y axis represents the tumour fraction D. Stacked bar plot representing CMS classification in WT p53 and p53 R273H COAD cohort (based on CMS classifier). Patients with R273H show higher proportion of CMS2 subtype(blue) and lower CMS1 subtype(yellow). The Y axis represents the tumour fraction. E. Stacked bar plot representing CMS classification in WT p53 and p53 R282W COAD patients based on CMS classifier. Patients with R282W shows higher proportion of CMS4 subtype(green) and lower CMS1 subtype(yellow). The Y axis represents the tumour fraction. F. Box plot representing the distribution of CMS1 gene expression in p53WT (grey) and p53mut (red). All three different p53 mutant tumours associate with significantly lower expression of CMS1 genes. G. Box plot representing the distribution of CMS4 gene expression in p53WT (grey) and p53mut (red). R273H and R282W tumours associate with significantly higher expression of CMS4 genes.

### p53 missense mutations show distinct proportion of CMS subtypes and associated gene expression

CRCs have been divided into four distinct consensus molecular subtypes (CMS) based on epigenomic and transcriptomic signatures [29]. The novel CMS classification system categorizes CRCs into four distinct subtypes: CMS1 (microsatellite instability [MSI]-immune), CMS2 (canonical), CMS3 (metabolic) and CMS4 (mesenchymal) [29]. Each of these CMS subtypes has a characteristic molecular background. The subtype has been demonstrated to be a prognostic factor. Initial classification has shown an association of Tp53 mutations with more CMS2 and CMS4 subtypes [30], however the variation in CMS subtypes with distinct mutational status of p53 has not investigated. To understand the subtypes associated with each of these three different p53 mutations, we classified each mutation types into different subtypes using their gene expression profile and identified different proportions of CMS subtype to be associated with distinct p53 mutations (Fig 2C-E, Fig S3A-C). Of these, the R282W mutations showed the highest fraction of CMS4 subtype and R273H, the highest fraction of CMS2 subtype. These results point to distinct molecular signatures associated with different p53 mutations.

To further correlate and understand the expression changes of each CMS subtype with distinct p53 mutations, we correlated each of the CMS gene signatures with different p53 mutation types and observed CMS1 genes with higher expression in p53WT tumours (Fig 2F) while CMS4 gene signatures showed significant elevated expression in R273H and R282W mutations but not with R175 tumours (Fig 2G), suggesting stromal infiltration, higher metastasis and poor prognosis associated with these mutations. However, we observed a significant gain in expression profile of CMS2 and decrease in expression of CMS3 genes across all the three tumours (Fig S2D-E)

### Distinct Methylation profiles with different missense p53 mutation in colon cancer

To understand the epigenetic changes associated with the p53 mutations under study, we explored the methylation profiles of different p53 mutation in colon tumours. WTp53 is known to suppress the expression of the *de novo* DNA methyl transferases DNMT3A and DNMT3B while upregulating Tet1 and Tet2, which promote DNA demethylation. In mouse embryonic fibroblasts, loss/mutation of p53 has been shown to upregulate DNMT3A [31].

Although direct association of DNA methylation with WTp53 has been investigated, the alterations in DNA methylation with respect to different hotspot mutation of Tp53 have not been explored. Towards this, we first explored the expression profile of DNMT3A in colon cancer patients harboring the three distinct mutations of Tp53. We observed significant elevation in the levels of DNMT3A in two of the three mutations under study, namely, the R175H and R273H mutations (Fig S3A).

We also observed distinct methylation profiles with each p53 mutation subtypes (Fig 3A-C). The R273H mutation exhibited significant hypermethylation while mutations R282W and R175 show significant hypomethylation profiles across patients. A total of 78,278, 50,363 and 30,240 probes were found to be significantly differentially methylated in R175H, R273H and R282W respectively. The density distribution profile confirms the association of hypermethylation with and hypermethylation with regard to R175H and R282W mutations (Fig3A-C). We also examined the distribution of the differentially methylated CpGs (DMCs) in various functional genomic regions such as promoters, CpG islands (CGIs), and CGI promoters (Fig 3D-F and Fig S3C-E).

**Fig 3:**
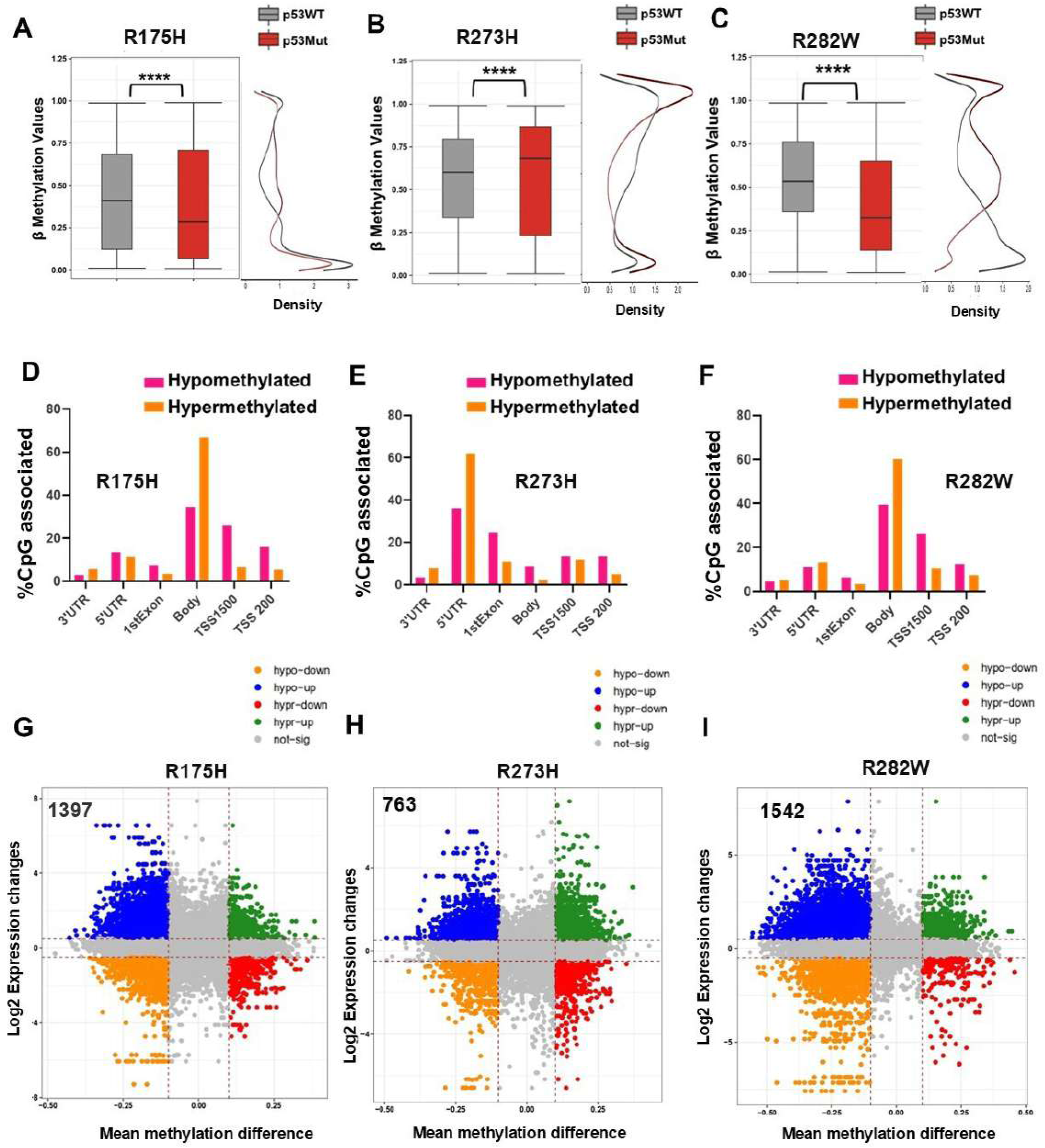
Distinct Methylation profiles with different missense p53 mutation in colon cancer. A. Box and density plots representing distribution of methylation values (β values) of significant probes (n=78,278) in Tp53 WT (grey) and Tp53R175H tumours (red). Tumours with R175H mutation show significantly lower methylation values as compared to p53 WT tumours. B. Box and density plots representing distribution of methylation values in Tp53 WT (grey) and Tp53R273H tumours (red) of significant probes(n=50,363). Tumours with R273H mutation show significantly higher methylation values as compared to p53 WT tumours. C. Box and density plots representing distribution of methylation values in Tp53 WT (grey) and Tp53R282W tumours (red)(n=30,240). Tumours with R282W mutation shows lower methylation values as compared to p53 WT tumours. D. Column bar chart representing distribution of hypermethylated(orange) and hypomethylated (pink) genomic regions upon R175H mutation. Hypermethylated regions are found distributed predominantly at gene bodies. E. Column bar chart representing distribution of hypermethylated(orange) and hypomethylated (pink) genomic regions upon R273 mutation. Hypermethylated regions mostly are distributed at 5’UTR. F. Column bar chart representing distribution of hypermethylated(orange) and hypomethylated (pink) at genomic regions with R282W mutation. Hypermethylated regions mostly are distributed at gene body. G. Integration of transcriptome and methylome data of colon tumours harbouring R175H mutation in Tp53. Starburst plot integrating differential DNA methylation and gene expression analysis. Genes that are hypermethylated and down-regulated genes are indicated in red; hypomethylated and up-regulated genes in blue; hypermethylated and up-regulated genes in green; or hypomethylated and down-regulated genes in orange. Hypomethylated genes correlate with increase in expression profile of genes in patients harbouring R175H tumours. X axis represents the DNA methylation changes (mean difference of β-values, Δβ) while the Y axis represents the gene expression changes (Log2 Fold Change) H. Integration of transcriptome and methylome data of colon tumours harbouring R273H mutation in Tp53. Starburst plot integrating differential DNA methylation and gene expression analysis. Hypermethylated and down-regulated genes are seen in red; hypomethylated and up-regulated genes in blue; hypermethylated and up-regulated genes in green; or hypomethylated and down-regulated genes in orange. Hypomethylated genes correlate with increase in expression profile of genes with R273H tumours. X axis represents the DNA methylation changes (mean difference of β-values, Δβ) while the Y axis represents the gene expression changes (Log2 Fold Change). I. Integration of transcriptome and methylome data of colon tumours harbouring R282W mutation in Tp53. Starburst plot integrating differential DNA methylation and gene expression analysis. Indicated are genes that are hypermethylated and down-regulated genes (red); hypomethylated and up-regulated genes (blue); hypermethylated and up-regulated genes (green); or hypomethylated and down-regulated genes (orange). Hypomethylated genes correlates with increase in expression profile with R282W tumours. X axis represents the DNA methylation changes (mean difference of β-values, Δβ) while the Y axis represents the gene expression changes (Log2 Fold Change).

We found that the open sea regions were hypermethylated while CpG islands were mostly hypomethylated irrespective of the type of p53 mutation. Further dissecting the distribution of DMCs across gene regions (TSS1500, TSS200, 5′ UTRs, first exons, gene bodies, and 3′ UTRs) we observed R273H mutation hypermethylated regions enriched at 5’UTR while R175H and R282W hypermethylated sites were enriched at gene body. These observations suggest that R175H and R282W follow a similar DNA hypo methylation profile while R273H follows a distinct hypermethylation profile.

One of the three major pathways associated with CRC pathogenesis is the CpG island methylator phenotype CIMP [32]. The CIMP subtypes are characterized by eight markers (CACNA1G, p16CDKN2A, CRABP1, IGF2, hMLH1, NEUROG1, RUNX3, and SOCS1) which are used to classify CRC subgroups into three different CIMP subtypes. If 1 to 5 out of the 8 markers show promoter methylation, it is classified as CIMP-low, when none of the markers is methylated, it refers to CIMP-0. If 6 to 8 out of 8 markers have promoters’ methylation, it is classified as CIMP-high. Colon cancer patients with CIMP-high CRC are known to showed a trend of decreased cancer-specific survival compared with CIMP-negative [33]. However, only about 11% of Tp53 mutations are associated with CIMP(+ve) phenotype. To understand the methylation profiles of these CIMP markers with these distinct p53 mutations, we compared the methylation values of these 8 markers with p53 WT tumours and observed that all of these CIMP markers are hypomethylated in each of the three mutations, suggesting that these p53 missense mutations are characterized with low CIMP phenotype (Fig S3B). This observation further correlates with epigenetic and genetic stratification of colon cancer cell lines, where TP53 mutation correlates with CIMP negative phenotype [34].

### Hypomethylated EMT-TFs correlate with increased gene expression on R273H mutation

In order to understand the functional relevance of hyper and hypomethylated DNA regions in each of the p53 mutations considered, we integrated the differentially methylated profiles obtained from these patients with changes in their gene expression profile. The genes were classified into four groups based on the intersection between the DMGs and DEGs: hypermethylated–upregulated (hyper–up), hypermethylated–downregulated (hyper–down), hypomethylated–upregulated (hypo–up), and hypomethylated–downregulated (hypo–down) genes (Fig 3G-I). We observed a total of 158 common genes hypomethylated and upregulated among all the three different p53 mutations. We observed zinc finger E-box binding homeobox (ZEB1) to be hypomethylated and upregulated with all the three different p53 mutations.

ZEB1 belongs to the EMT-zinc finger transcription factor family and is involved in crucial mechanisms including tumor progression [35]. Recent work has observed a frequent ZEB1 hypermethylation in CMS1 patients and this hypermethylation of ZEB1 is a good prognostic factor related to disease-free survival and overall survival in colon cancer. Interestingly, we also observed a unique hypomethylated signature of EMT-TFs VIM and SNAI2 with R273H mutation. SNAI2 is a key regulator of epithelial-to-mesenchymal transition (EMT) and its aberrant expression has been observed in various cancer types and predicts poor prognosis in cancer patients [36]. We observed a distinct hypomethylation of key EMT-TFs VIM and SNAI2 which correlates with increased expression exclusively with R273H suggesting that DNA methylation plays an important role in regulating R273H mutation in colon tumours (Fig 4B, E). In order to understand the functional relevance of these hypomethylated upregulated genes with distinct p53 mutation, we performed single sample gene set enrichment analysis (ssGSEA) and observed significantly upregulated TGF-β signalling (Fig 4G) and EMT-pathways (Fig 4H) to be enriched with R273H mutation, suggesting that the metastasis exhibited by the p53 R273H mutation is affected through DNA methylation in colon tumours. The hypomethylated regions upon R273H mutation also correspond to the increased expression of genes such as RUNX2 and TEAD1 involved in the YAP/TAZ signalling pathway (Fig S3F-G) in R273H colon tumours, suggesting a novel epigenetic regulation of genes involved in colon cancer metastasis. This highlights the importance of devising epigenetic based therapy based on Tp53 mutation status.

**Fig 4:**
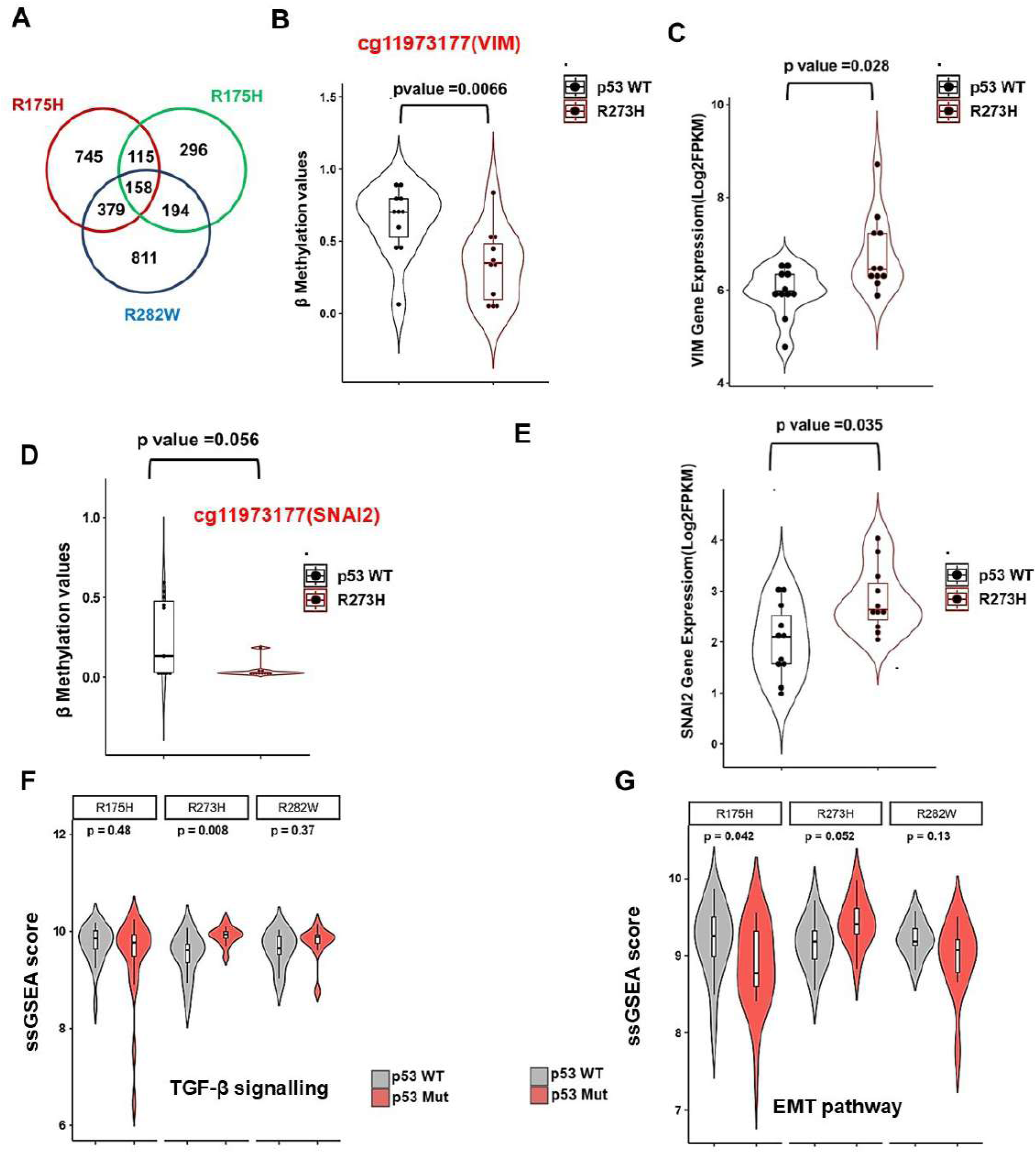
Hypomethylated EMT-TFs correlate with increased gene expression profiles in patients with R273H mutation. A. Venn diagram representing unique and common upregulated hypomethylated genes with different p53 mutation as obtained by integration of methylome and transcriptome data. R273H shows unique EMT-TF signatures with hypomethylated promoters correlates with upregulation of key EMT-TFs including VIM and SNAI2. B. Box and Violin plot representing methylation levels of VIM (cg11973177) probes in p53 WT (black) and R273H mutation (red). R273H mutation shows hypomethylation a VIM promoters. C. Box and violin plots representing vimentin (VIM) expression profile in p53 WT (grey) and R273H tumours. R273H shows significant higher expression of VIM as compared to p53WT tumours. D. Box and Violin plot representing the distribution of methylation levels of cg11973177(SNAI2) probes in p53 WT (black) and R273H mutation(red). R273H mutation shows hypomethylation at SNAI2 promoters. E. Box and violin plot representing SNAI2 expression profile in p53 WT (grey) and R273H tumours. R273H tumours show significantly higher expression of SNAI2 as compared to p53WT tumours. F. Box and Violin plot representing ssGSEA score of hypomethylated upregulated genes in p53WT (grey) and p53 mutant (red) tumours. R273H mutation correlates with significantly higher ssGSEA score involved in TGF β signalling pathway as compared to R175H and R282W tumours implying the involvement of EMT and metastasis. G. Box and Violin plot representing ssGSEA score of hypomethylated upregulated genes in p53WT (grey) and p53 mutant (red) tumours. R273H mutation correlates with significantly higher ssGSEA score involved in EMT pathway as compared to R175H and R282W tumours.

**Fig 5.**
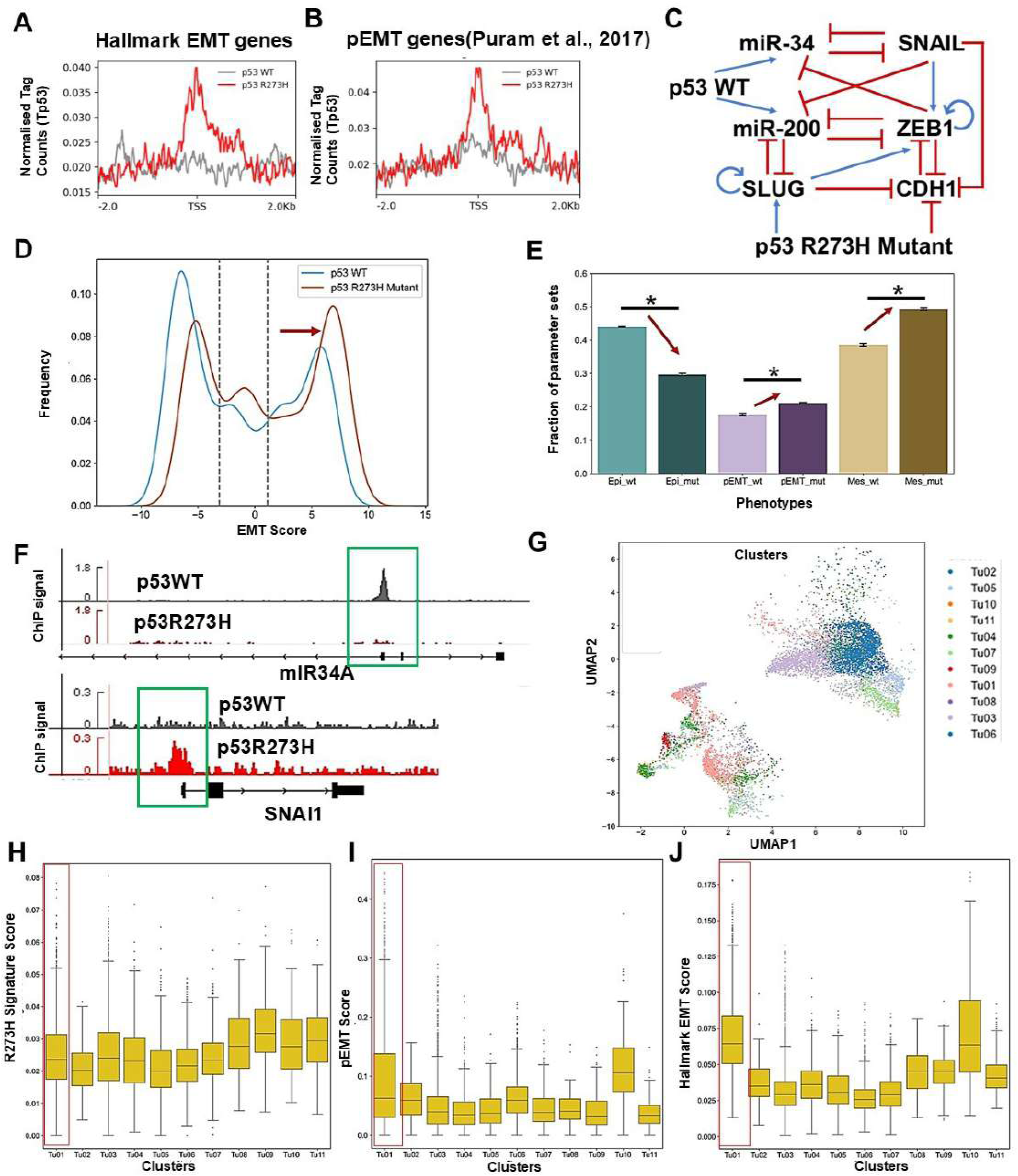
Mutant p53 R273H reshapes GRN dynamics and enriches at partial/mesenchymal EMT states in colorectal cancer. A. PlotProfile representing WTp53 signal (grey) and Tp53 R273H signal (red) at the Hallmark EMT genes (+/-2kb of TSS). Tp53R273H signal corresponds to a higher signal intensity as compared to that of p53WT at the promoters of hallmark EMT genes. B. PlotProfile representing WTp53 signal (grey) and Tp53 R273H signal (red) at the pEMT genes (+/-2kb of TSS). Tp53R273H signal corresponds to a higher signal intensity as compared to that of p53WT at the promoters of pEMT genes. C. **In silico GRN modeling and phenotype quantification.** Schematic of the gene regulatory network (GRN) incorporating key EMT regulators and p53 nodes. Red hammerheads represent inhibitory interactions; blue arrows indicate activatory links. D. Kernel density estimate (KDE) plots show the distribution of EMT scores under wild-type and R273H mutant p53 overexpression. The red arrow indicates a rightward shift in EMT score distribution for the mutant, suggesting increased mesenchymal bias. E. Bar plots display the fraction of steady states classified as epithelial, partial EMT, or mesenchymal across triplicate simulations of the GRN under p53 WT and R273H mutant overexpression. Arrows indicate trends in phenotype redistribution, with mutant p53 favouring hybrid and mesenchymal states. F. IGV screenshot representing the alterations in the levels of p53WT(HCT116)(grey) and HT29 (p53R273H)(red) CRC cells at mIR34A loci and SNAI1. P53WT enriched at the promoters of miR34A while p53R273H enriched at the promoters of SNAI1. G. Single-cell transcriptional heterogeneity in CRC: UMAP projection of tumour cells (GSM7058755) reveals ten transcriptionally distinct tumour clusters. H. **H-J** Distribution of enrichment scores across tumour subpopulations: Box plots showing signature scores for (H) R273H gene signatures (I) pEMT (J) Hallmark EMT gene signatures. The red box highlight a representative cluster (e.g., Tu1) where R273H enrichment co-occurs with elevated partial and full EMT programs, indicating mutant p53-associated phenotypic reprogramming.

### Mutant p53 R273H alters the EMT landscape by promoting phenotypic plasticity through reprogramming of core gene regulatory network in colorectal cancer

To understand the regulation of mutant R273H on these differentially methylated genes and to compare the direct and indirect regulation of Tp53 on genes responsible for EMT transition and partial EMT genes, we performed genome wide profiling using chromatin immunoprecipitation followed by sequencing (ChIP-Seq) of Tp53 in HT29 (p53R273H) and HCT116 (p53WT) CRC cell lines (GSE86164). We identified regions enriched with p53WT (Fig S4A) and p53R273H mutation binding sites (Fig S4B) and observed a significantly higher enrichment of R273H at the promoters of hallmark EMT genes [24] (Fig 4A, Fig S4C) and partial EMT genes [28] as compared to p53WT (Fig 4B, Fig S4D) suggesting a direct influence of R273H in regulation of partial EMT and mesenchymal states. Mutations in TP53, particularly the R273H hotspot mutation, are known to profoundly alter cellular phenotypes by modulating gene regulatory networks that govern cell fate decisions. The R273H mutation in TP53 reprograms colorectal cancer (CRC) cells along the epithelial-mesenchymal transition (EMT) axis, favouring hybrid and mesenchymal phenotypes. This shift reflects enhanced phenotypic plasticity, a hallmark of tumour progression, invasion, and therapy resistance. Mutation at the R273 codon of p53 not only abrogates its tumour-suppressive functions but also imparts gain-of-function properties that reshape the regulatory landscape governing cell state transitions. To investigate the molecular underpinnings of these phenotypic shifts, we constructed a gene regulatory network (GRN) that integrates experimentally validated interactions and captures emergent dynamics of cell state plasticity. This GRN includes core EMT regulators such as SNAIL, SLUG, ZEB1, and CDH1; microRNAs like miR-34 and miR-200; and p53 in its wild-type and R273H mutant forms. The EMT circuitry is modelled as two interconnected modules: the miR-34/SNAIL and the miR-200/ZEB axes, as described in earlier work [37]. Additionally, interactions involving SLUG, ZEB1, CDH1, and miR-200 were integrated based on established literature [38]. Crucially, wild-type p53 activates miR-34 [39] and miR-200 [40], reinforcing the epithelial state, whereas the R273H mutant disrupts this balance by activating SLUG (S. P. Wang et al., 2009,J. Kim et al., 2014) and repressing CDH1 [43], thereby favouring mesenchymal traits (Fig 4C). To simulate the cell-state dynamics governed by the GRN, we employed Random Circuit Perturbation (RACIPE) [44] framework, a computational method that captures the intrinsic variability and emergent behaviour of regulatory networks. RACIPE models the GRN as a system of coupled ordinary differential equations (ODEs), assigning one ODE per network node, with kinetic parameters for production, degradation, and regulatory interactions sampled from biologically plausible ranges. By solving these ODEs across thousands of randomly sampled parameters sets and initial conditions, RACIPE generates a robust ensemble of steady-state solutions. These steady states collectively define the “possibility space” of phenotypes that can arise from the underlying network topology. We simulated this GRN under three scenarios: the baseline (no p53 perturbation), wild-type p53 overexpression (mimicking a homozygous wild-type scenario), and R273H mutant p53 overexpression (mimicking a homozygous mutant scenario). To quantify the EMT status of each steady state, we computed an EMT score (see Material and methods), which allowed us to map the phenotypic spectrum along the epithelial– mesenchymal axis. Kernel density estimates (KDEs) of the EMT score distributions revealed a rightward shift in the R273H mutant condition compared to wild-type p53, indicating a higher prevalence of mesenchymal-like and hybrid phenotypes (Figure 4D). This suggests that mutant p53, unlike its wild-type counterpart, biases the network dynamics towards more partial and mesenchymal cell states. To further dissect this shift, we stratified the steady-state population into epithelial, hybrid/partial EMT and mesenchymal phenotypes using EMT score cutoffs determined from local minima in the baseline EMT score distribution. Comparative analysis of phenotypic frequencies showed that the overexpression of p53 R273H led to a marked reduction in epithelial states and a concomitant increase in both hybrid and mesenchymal populations relative to wild-type p53 (Figure 4E).

We next identified and validated enrichment of p53WT and p53 R273H at specific epithelial, mesenchymal and pEMT genes. We identified enrichment of p53WT at the promoter of miR34A but not R273H (Fig 4F), suggesting that wild-type p53 activates miR-34, reinforcing the epithelial state, whereas the R273H mutant shows higher enrichment at SNAI1(EMT-TF) (Fig 4F) and THBS1 (pEMT) gene (Fig S4E) as compared to p53WT suggesting the differential regulation of epithelial and mesenchymal markers, disruption of which favors mesenchymal traits. These findings underscore the role of mutant p53 in modulating the phenotypic landscape accessible to cancer cells, enhancing cellular plasticity and potentially contributing to metastasis and therapeutic resistance.

### Single-cell transcriptomic analysis reveals R273H-linked enrichment of partial and mesenchymal EMT phenotypes across tumour subpopulations in CRC

To assess the transcriptional impact of the p53 R273H mutation, we analysed single-cell RNA sequencing data from colorectal cancer (CRC) tumour tissues (GSE225857; sample GSM7058755). UMAP-based dimensionality reduction of ten distinct tumour cell clusters, reflected substantial intra-tumoral heterogeneity (Figure 4G). We computed enrichment scores for a previously curated R273H-associated transcriptional signature [10] to identify cells transcriptionally influenced by this mutation. These scores were mapped onto the UMAP projection, revealing discrete subpopulations enriched for the R273H program (Fig S4F). To further evaluate the EMT landscape, we assessed enrichment of both partial EMT (pEMT) [28] and Hallmark EMT signatures [24] across the same single-cell dataset (Fig SFG, Fig S4G). Quantitative analysis of these pathway scores across the ten tumour clusters, showed clusters with elevated R273H signature enrichment (Fig 4H), particularly Tu1, also showed high levels of both pEMT (Fig 4I) and hallmark EMT scores (Figures 4J). This correlation suggests that the transcriptional program associated with mutant p53 R273H preferentially aligns with partial and mesenchymal-like EMT states.

## Discussions

The high prevalence of Gain of Function (GoF) mutations, particularly missense/hotspot mutations across various cancer types suggests a selective advantage during cancer progression [45,46, 20]. The broad mutation spectrum of p53, most of which lies in the DNA binding domain of Tp53 varies across types of malignancies and associates with varied phenotypes [10,12].

More than 50% of Tp53 mutations are associated with metastatic colorectal cancer and different GoF mutations have very different clinical outcomes [47], thus making it crucial to understand the phenotype and regulatory mechanisms associated with different Tp53 mutations. In this work, we have attempted to dissect and unravel the transcriptomic and epigenomic signatures associated with three most prevalent and distinct hotspot mutations in Tp53 in patient colon tumours through TCGA COAD.

Exploratory analysis through TCGA-COAD project identified three distinct and prevalent mutations in Tp53 viz., R175H (N=25), R273H (N=13) and R282W (N=11). These distinct mutations show varied survivability profiles and unique transcriptomic signatures. Colon cancer patients with R273H mutation were found to be associated with lower survival rates as against R175H and R282W mutations of p53. Gene set Enrichment (GSEA) analysis and single sample gene set enrichment analysis (ssGSEA) analysis further reveal unique reactome signatures, with R273H mutation involved in YAP/TAZ signalling with an upregulation of YAP1 and TEAD1 genes implying a vital role for the p53R273H mutation in cancer stemness, renewal and tumour initiation in colorectal cancer mostly by modulating the expression of genes involved in Hippo signalling pathway. The transcriptomic profiles for mutation R175H show enrichment in WDR5 containing histone modifying enzymes, The enrichment of WDR5 containing histone modifying further reveals an epigenetic pathway associated with R175H mutation in colon cancer. CMS classification on each of the different p53 mutant tumours reveals a very distinct enrichment of CMS4 subpopulation with R282W mutation. We identified R273H and R282W mutation to be associated with significantly higher expression of CMS4 gene signatures as compared to R175H tumours signifying higher metastatic and cancer metastasis properties with R273H and R282W as compared to R175H tumours.

A significant upregulation of DNMT3A was also seen in R175H and R273H tumours but not with R282W mutation tumours. TP53 in its wild type form functions as a direct repressor for DNA methyl transferases including DNMT3A and DNMT3B [31], which correlates with increase in expression profile observed upon R273H and R175H mutation. Aberrant DNA methylation is the most extensively characterized epigenetic alteration observed across all stages of colorectal cancer progression. However, the methylation patterns associated with different p53 mutations have not been explored so far.

Investigations on the DNA methylation profile of each of these three distinct p53 tumours revealed a distinct methylation pattern associated with different Tp53 mutation. These novel DNA methylation signatures associated with different p53 mutations further signifies an importance to understand the type of Tp53 mutation and the phenotype associated with each tumours. Interestingly, we observed R273H tumours to have higher methylation profiles (Hypermethylation), while R173H and R282W tumours to have lower methylation values (Hypomethylation) in colon tumours reflecting a distinct methylation signature across different p53 tumours. This led us to hypothesise that WTp53 tumours in general, repress the DNA methyl transferases, and upon acquiring gain of function mutations lead to a differential regulation of DNA methyl transferases establishing a very distinct methylation profile. DNA methylation markers have been proposed as potential biomarkers for early colorectal cancer progression.

Functional integration of DNA methylation profile with transcriptomic profile further identified unique EMT-TF signatures to be associated with R273H tumours. We observed that ZEB1, an EMT-TF is hypomethylated and upregulated across all the three tumour classes, while Vimentin (VIM) and SNA12 are hypomethylated and upregulated only with R273H tumours highlighting a novel epigenetic circuit driving cancer towards metastasis with R273H tumours. Aberrant DNA methylation of VIM in fecal DNA was proposed in detecting CRC diagnosis with an overall specifity of 88% suggesting that these methylation pattern around key EMT-TFs can be stratified and included as a biomarker for detection of CRC pathogenesis [47,48]. Single sample gene set enrichment analysis on the hypomethylated upregulated genes revealed that R273H tumours are significantly enriched with higher TGFβ signalling and Epithelial to Mesenchymal transcription (EMT) pathways highlighting the importance of different combinatorial therapeutic regime depending upon the p53 mutation status.

Although various studies have highlighted the interaction of YAP1 and Tp53 mutation in controlling the expression of various genes involved in tumour progression [49] using breast cancer model system, the epigenetic circuitry or methylome signatures associated with YAP/TAZ signalling pathway with R273H has not been previously investigated. This work hence provides novel insights into unique epigenomic and transcriptomic signatures associated with the three prevalent missense mutations of Tp53 in colon cancer.

Epithelial to mesenchymal transition in cancer cells are characteristic of dynamic and reversible transitions between epithelial and mesenchymal phenotypes to endure the environmental stress and increase the chances of successful metastasis. EMT spectrum is comprised of several intermediate cell states collectively termed hybrid epithelial-mesenchymal (hybrid E/M) characterized by both epithelial markers (E-cadherin, cytokeratins, claudins, occludins) and mesenchymal markers (N-cadherin, vimentin, fibronectins), and they may display phenotypic characteristics of both cell types [50, 23]. However, alteration in transcriptional landscape of key EMT and pEMT genes with respect to different GoF mutation status of Tp53 has not been well established. Our analysis reveals that R273H associates with elevated mesenchymal and pEMT gene signatures. Further analysis of ChIP seq data reveals that R273H mutation shows higher enrichment at the promoters of EMT signatures and pEMT thus influencing the pEMT and mesenchymal states. The gene regulatory network further establishes that overexpression of R273H, unlike its wild-type counterpart, biases the network dynamics towards more partial and mesenchymal cell states by reducing the epithelial states and a concomitant increase in both hybrid and mesenchymal populations relative to wild-type p53. Single cell transcriptomic analysis of colon tumours further validates that R273H gene signatures correlate with pEMT and EMT like sates. This correlation suggests that the transcriptional program associated with mutant p53 R273H preferentially aligns with partial and mesenchymal-like EMT states.

Epigenetic regulation through DNA methylation/demethylation plays an important role in regulating hybrid E/M conditions [51]. DNMT1 in particular has demonstrated an important role in driving hybrid E/M states thorough its interaction with Snail1/2 [52, 53, 54]. Interestingly. we observed a significant hypomethylation profile of SNAI2 and VIM key EMT-TF upon R273H mutation.

These observations also highlight the importance of stratifying tumours based on the type Tp53 mutation to design novel combinatorial therapeutic regime. Methylation based therapies including AZA, DAC, SGI-110, or TMZ are currently in clinical trials for treatment of colorectal cancer patients [55]. Pan-DNMT inhibition has shown to prevent TGF-β-induced hybrid E/M gene expression and metastatic phenotypes in ovarian and prostate cancer cells [53,56]. However, new stratification-based studies including CMS subtype and mutational status may reveal further exploration and strategic combination for devising new therapeutic regimen for colorectal cancer treatment. This work reports a hitherto unexplained relationship between p53 mutant specific epigenetic signatures and partial EMT whose novelty shall help design targeted epigenetic therapies for p53 mutant tumours.

## Material and Methods

### Identification of mutant p53 patient sequencing data

Primary colon tumours with transcriptomic profiling data were chosen from the TCGA -COAD (Colon Adenocarcinoma). The samples of patients which harboured three prevalent gain of function (GoF) missense mutations in the DNA binding domain of (R175H, R273H and R282W) were chosen for the analysis. Somatic mutations of these patients available in the Mutation Annotation Format (MAF) were retrieved for these samples. Maf analysis was done using the “maftools” R package [57] for visualization and summarization of MAF files. The tumours were stratified on the basis of the presence of single and distinct p53 missense mutations into three distinct subtypes (a) R175H (N=25) (b) R273H (N=13) and (c) R282W (N=11). Samples without any mutation in p53 were considered p53 WT tumours.

### Differential Gene Expression Analysis

The raw expression counts and TPM count of genes for distinct p53 mutational type (a) R175H (N=25) and p53WT (N=25) (b) R273H (N=13) and p53WT (N=13) and (c) R282W (N=11) and p53WT (N=11) tumour samples were obtained from the GDC portal using TCGAbiolinks [58]. Further, the count matrix was normalized (using TCGAanalyze_Normalization function) and genes with low expression count were filtered out (using TCGAanalyze_Filtering with method = quantile, and qnt.cut = 0.25). Differential expression analysis was carried out using edgeR through TCGAanalyze_DEA function (with method = glmLRT). Genes with |log2foldchange| > 0.5 and pvalue <= 0.05 were defined as significantly differentially expressed.

### CMS Classification

In order to classify the different samples into their CMS subtypes, gene expression counts were used. Ensembl IDs were translated to Entrez IDs with the biomart Bioconductor package [59]. Raw counts were log transformed and batch effects were removed from sequencing platforms before processing with CMS classification. The random forest models called CMS classifier [29] and CMS Caller [60] were applied on the identified p53 WT patients and p53 missense mutation cohorts through (TCGA-COAD). CMS class was assigned according to the default settings (minCor = 0.15, minDelta = 0.06). Box plot representing gene expression profiles of different CMS subtypes in each mutation cohort were plotted using ggplot2 in R. Student-t test was used to evaluate the significance in expression profiles for each of the mutations.

### Differential methylation analysis

Illumina HumanMethylation450K array data from stratified colon tumours based on p53 mutational status (A) R175H (N=16) and p53 WT (N=16) (B). R273H (N=11) and p53 WT (N=11) (C) R282W tumours (N=7) and p53WT(N=7) were obtained from TCGA using TCGAbiolinks [58]. The methylation levels as beta values (β), defined as the ratio of the intensities of methylated and unmethylated alleles were used for further analysis. Differential methylation analysis was performed between individual missense and WT tumor samples. Probes containing SNPs, probes in chromosome X, and probes with more than 10% missing values were excluded from the analysis. The Wilcoxon rank-sum test was used to determine the differentially methylated CpGs (DMCs). DMCs were reported as those probes which showed the mean methylation difference >0.1 with a p-value cut off less than 0.05. The probes were annotated by using the Bioconductor package with the human genome assembly GRCh37.

### Gene Set Enrichment Analysis (GSEA) and single set gene set enrichment analysis (ssGSEA)

To identify the significantly enriched biological processes or pathways associated with a set of differentially expressed genes (pval<0.05 and log2FC|0.5|), GSEA and ssGSEA were performed using GSEA [24]. Gene set c2.cp.reactome.v2024.1.Hs.symbols.gmt and hallmark gene sets were selected for the enrichment analysis using the default settings and statistically significant enrichment results were determined based on the p-value threshold of 0.05.

### Integration of DNA methylation and gene expression data

The differentially methylated regions and the differentially expressed genes were integrated to identify genes which are functionally different. Genes which were hypermethylated– downregulated (hyper–down) and hypomethylated–upregulated (hypo–up) with a logFC|0.5| and methylation difference delta value of |0.1| (Fig4A-C) and p-value of <0.05 were defined as functionally relevant genes. GSEA and ssGSEA was performed for these functionally relevant genes against hallmark gene sets using GSEA [24]. Box plot representing ssGSEA score was plotted using ggplot2 in R. Wilcoxon Rank Test was used to evaluate the significance.

### Survival Analysis

Kaplan-Meier (KM) analysis was conducted to evaluate the prognosis of the tumours based on the mutation status of Tp53. Clinical data and mutational status of Tp53 were derived from TCGA for which transcriptome profile was available. Cohort was divided into two groups p53WT and p53 missense mutation. Further each missense mutation was divided into three different types. KM analysis was conducted using ggsurvfit (1.1.0) and p values were calculated using the log-rank test in R.

### Cell Culture

Colon adenocarcinoma cell line HT29 harbouring p53R273H mutation was procured from NCCS, Pune and was maintained in Dulbecco’s Modified Eagle’s Medium and Hams-F12 (DMEM-F12) (#12500062 Invitrogen-Gibco) supplemented with 10% FBS (#16000044 Invitrogen-Gibco) and 1% penicillin-streptomycin (#1507070063 Invitrogen-Gibco). The cells were maintained in humidified environment at 37°C in presence of 5% CO2.

### Chromatin immuno Precipitation assay and library preparation

2.5 million cells were seeded and cultured to attain 80-90% confluency in 100mm dishes. The cells were crosslinked with 1% formaldehyde (#219404790 MP BIO) at room temperature for 10 mins with constant shaking. Glycine was added to a final concentration of 125 mM to quench the formaldehyde for 5 mins. Cells were washed thrice with 1X ice cold PBS. Cells were scraped in 3 ml of 1X PBS and pelleted down at 2K rpm for 5 mins at 4°C and kept at −80° C. Nuclear lysis buffer (L2) supplemented with 1X PIC was added to the nuclear pellet and incubated on ice for 10 mins. Samples were sonicated using Covaris-S220. The cell lysate was cleared by centrifuging samples at 12K rpm for 12 mins. 100 ug of sheared chromatin was taken for each IP. Sheared chromatin was diluted by adding Dilution Buffer (DB) supplemented with 1X PIC in 1:1.5 (1 volume of sheared chromatin and 1.5 volumes of Dilution Buffer) ratio. 10% of diluted chromatin was set aside as Input. 2µl of antibody(p53(#7F5) Rabbit MAb)) was used for each IP. Dynabeads (Invitrogen-#140004D) were prepared by blocking in 1% BSA prepared in 1X PBS at 4°C for 1 hour followed by washing with 1X PBS. Immunoprecipitated DNA was collected by adding 14µl of BSA blocked beads to each sample and incubated at 4°C for 4 hours. Beads were collected, flow through was discarded and 800 µl of Wash Buffer I was added. Washings were carried out at 4°C on a rocking platform. Washings were sequentially repeated with Wash Buffer II, Wash Buffer III and 1X TE. Immunoprecipitated DNA was eluted by adding 200 μl of Elution Buffer for 45 mins at 37°C in a thermomixer with rpm of 1200. Eluate was transferred to fresh tubes and 14 µl of 5M NaCl was added and kept overnight at 65°C. Immunoprecipitated DNA was purified by Phenol:chloroform:isoamyl alcohol (Ambion-AM9732) method, followed by ethanol precipitation. The final dried DNA pellet was dissolved in 15µl of 1X TE. 1ng of the DNA was used for preparation of ChIP libraries using NEBNext® ltra™ II DNA Library Prep with Sample Purification Beads (#E7103L) and subjected to sequencing using illumina Hi Seq 2500 sequencing platform, 1x55 bp sequencing read length.

### ChIP-Seq data processing and analysis

Fast QC was done for all the samples to check for adapter content, PHRED score and GC content. Adapter content was removed using Trimmomatics (v 0.39). Trimmed reads were later mapped to the human genome (hg38) using Bowtie2(v 0.2.4.1) [61]. Sam files were converted into sorted and indexed bam files using samtools. Peak calling was performed using MACS2 [62] with a p-value threshold of 5E-10. For each condition, the corresponding input samples were used for background normalization. To compare the ChIP sequencing data sets in WT and p53 mutant cell line, the consensus set of peaks obtained in each condition were merged using the “merge” function from bedtools [63]. The density of reads in each merged region was quantified using normalized signal per million reads. The gained and lost sites were defined on the basis of normalized signal per million reads for which log2fold change (log2FC) was larger than >0.5. Profiles were obtained on a region of promoters of Hallmark EMT genes [24] and pEMT genes ([28]), and the average scores were plotted to generate averaged read density using Deeptools plotProfile [64].

### GRN simulations using RACIPE

To investigate the dynamic behaviour of the gene regulatory network (GRN), we employed the Random Circuit Perturbation (RACIPE) framework. RACIPE simulates the transcriptional dynamics of a regulatory network by translating its topology into a system of nonlinear ordinary differential equations (ODEs), where each gene or node is governed by rate-based expressions incorporating production, degradation, and regulatory influences. The framework generates an ensemble of models by randomly sampling biologically relevant kinetic parameters, such as production and degradation rates, Hill coefficients, and threshold levels, for each node and interaction. These parameters are varied across thousands of independent simulations, each run from multiple initial conditions, to capture the full range of potential steady-state behaviours dictated by the network structure.

For a node T in the network having Pi activating and Nj inhibiting nodes with incoming edges, the ODE generated by RACIPE to represent the time evolution of concentration of node T is as follows:

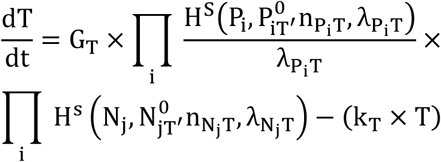

where the terms T, Pi, and Nj are concentrations of nodes at time t (gene expression levels), n is Hill coefficient showing the influence of Pi or Nj on T, λ is fold change in expression of T, caused by regulatory node Pi, or Nj, P_i_^0^ or N_j_^0^are threshold values of Hill function, GT and kT is production rate and degradation rate of node T respectively. H^S^ (shifted hill function) takes in the activatory/ inhibitory links in account to determine the production rate for the node and is defined by:

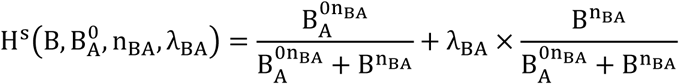

A key feature of RACIPE is the implementation of the “half-functional rule,” which estimates threshold parameters such that each regulatory interaction has approximately a 50% chance of being active across the ensemble. This ensures unbiased representation of all interactions and prevents the network topology from being effectively altered due to extreme parameter values. To facilitate comparison across nodes and conditions, steady-state expression values are Z-normalized based on the mean and standard deviation of the baseline circuitry for each node across all simulations. In this study, we simulated the GRN using 10,000 randomly sampled parameter sets per replicate, with 100 initial conditions per set, and performed simulations in triplicate to ensure robustness.

To quantify the epithelial–mesenchymal transition (EMT) spectrum from the simulated GRN steady states, we computed an EMT score based on the expression of key marker genes. The epithelial score was calculated as the sum of Z-normalized expression values of Cdh1, miR-200, and miR-34, while the mesenchymal score was derived similarly from Snail, Slug, and Zeb1. The final EMT score was defined as the difference between the mesenchymal and epithelial scores (EMT score = mesenchymal score − epithelial score), with higher values indicating a more mesenchymal-like state. This metric enabled classification of cell states along the epithelial– mesenchymal continuum.

### Analysis of Single cell Transcriptomic Data

Publicly available single-cell RNA-seq count matrix was obtained from the GEO dataset GSE225857, specifically focusing on non-immune tumour cells from sample GSM7058755. To assess the activity of specific pathways and gene signatures, including the p53 R273H signature, partial EMT (pEMT), and Hallmark EMT, we computed gene set enrichment scores using the AUCell algorithm [65], which estimates the relative activity of a gene set in each cell based on its expression ranking. These activity scores were used to map signature distributions across UMAP projections and tumour clusters.

## Data Availability

The somatic mutation profile, gene expression profile and methylome data used in study was obtained from https://portal.gdc.cancer.gov/. Other data generated in the study is available on request.

## Acknowledgements

VM was supported by ICMR, India (Grant Number 2020-5255/GENOMIC/ADHOC/BMS) and DST-SERB,I ndia (Grant Number CRG/2019/006509) for to VM

- MKJ was supported by Param Hansa Philanthropies
- HR was funded by the Council of Scientific and Industrial Research(CSIR), India
- The authors acknowledge the infrastructure support (HPC server) at IBAB funded by DBT Skill Vigyan Programme BT/HRD/01/012/2017 and the Centre for Disease Genomics, Big Data Analysis in Bioinformatics and Healthcare’ – BIC at IBAB funded by the DBT, India BT/PR40212/BTIS/137/40/2022
- We thank Dimple Notani for providing us the infrastructure for ChIP sequencing and for valuable discussions.

