## Supplementary figures and images for "Gain of Function p53 mutant R273H confers distinct methylation profiles and consequent partial or full EMT states to colon tumour"

### Supplementary Fig S1

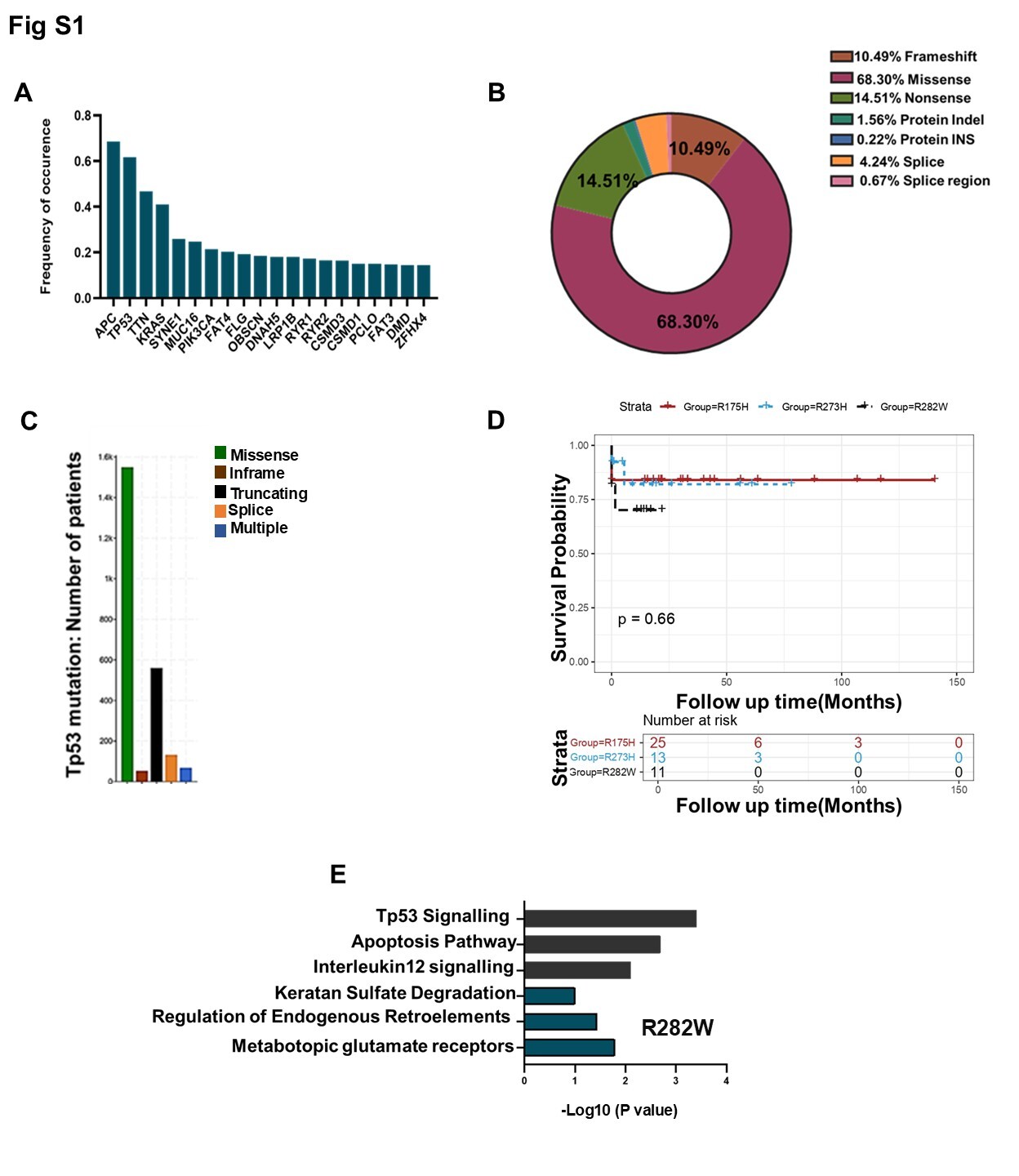

### Supplementary Fig S2

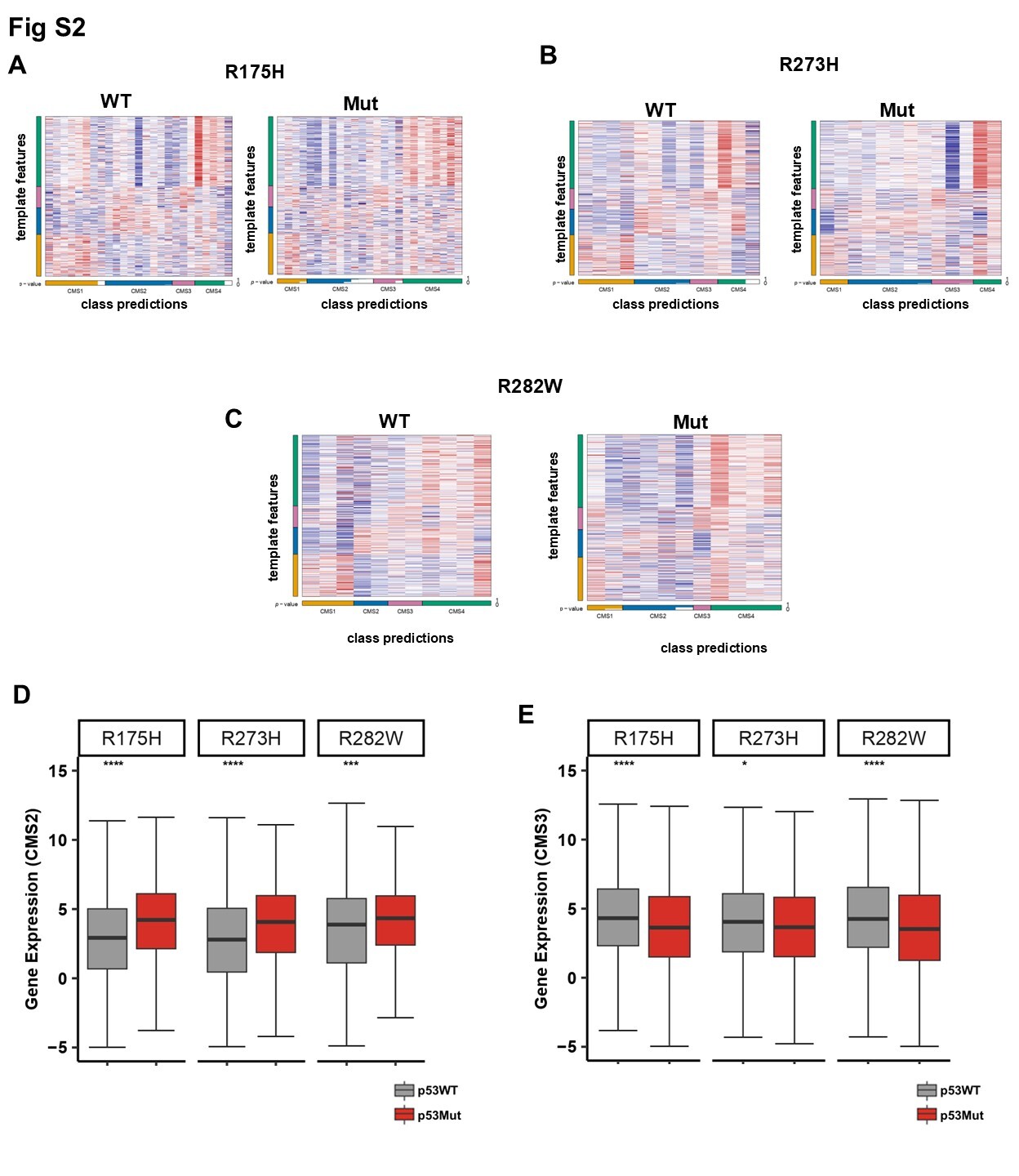

### Supplementary Fig S3

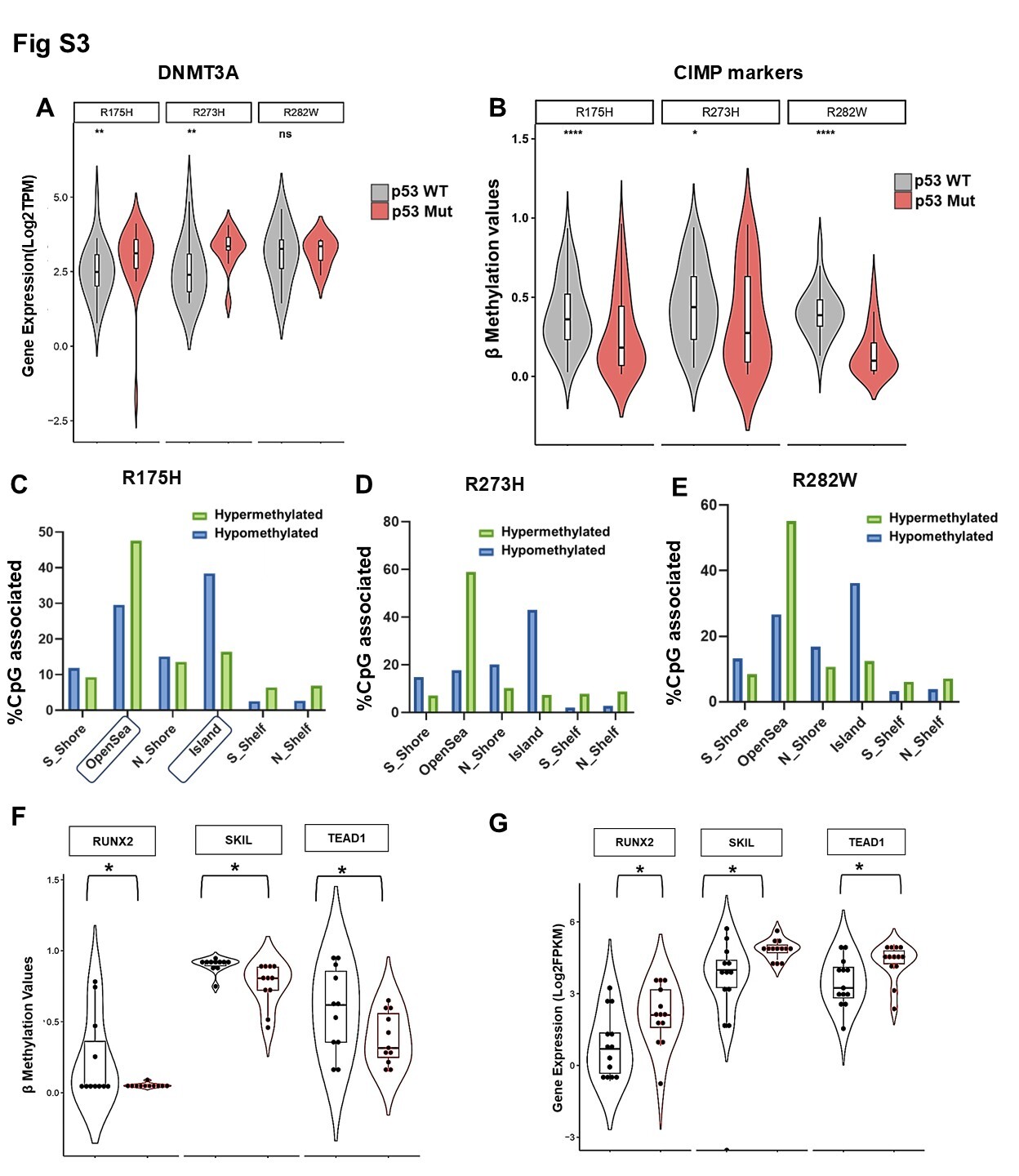

### Supplementary Fig S4

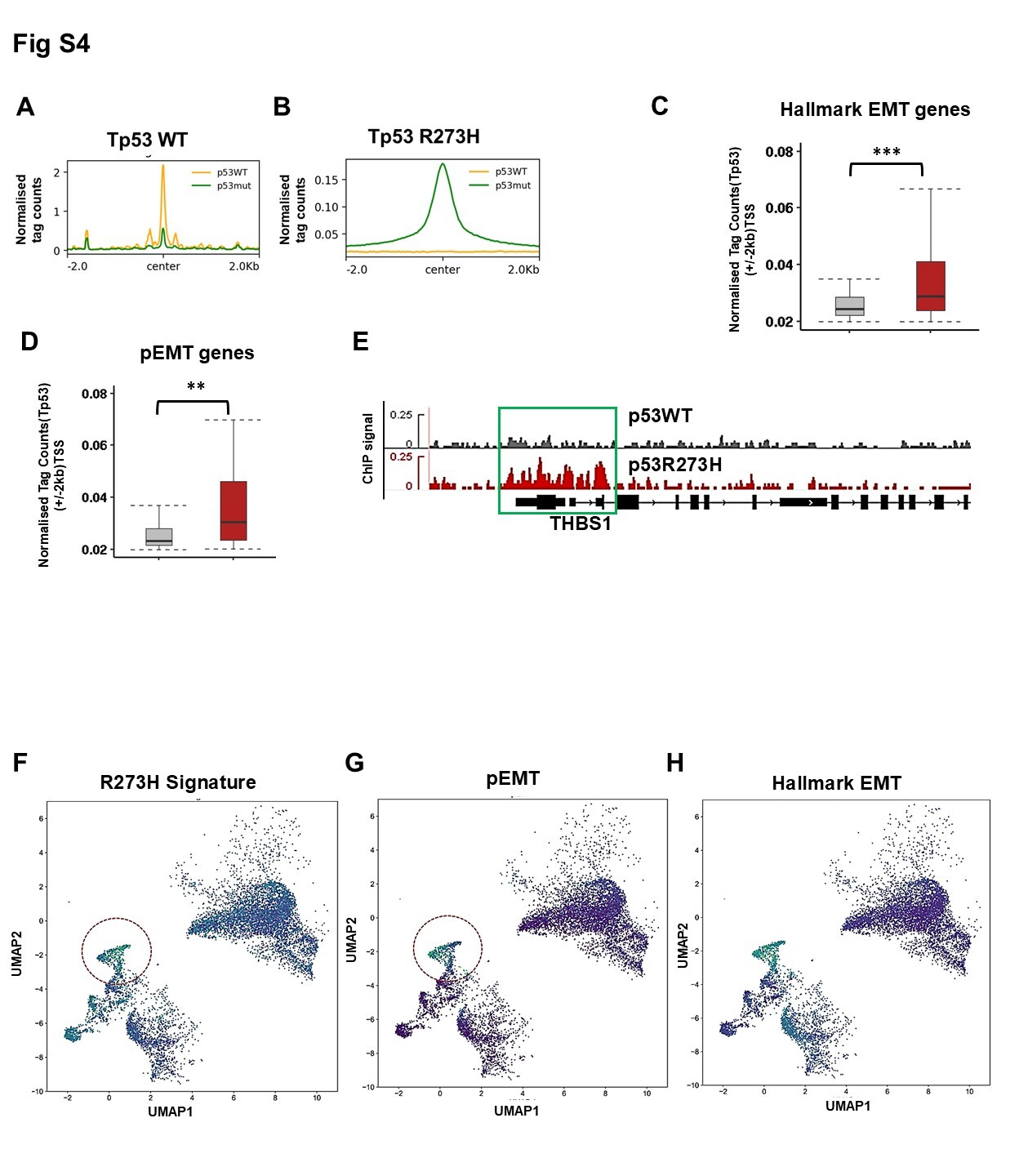
