## Supplementary Figure Legends for "Gain of Function p53 mutant R273H confers distinct methylation profiles and consequent partial or full EMT states to colon tumour"

Fig S1

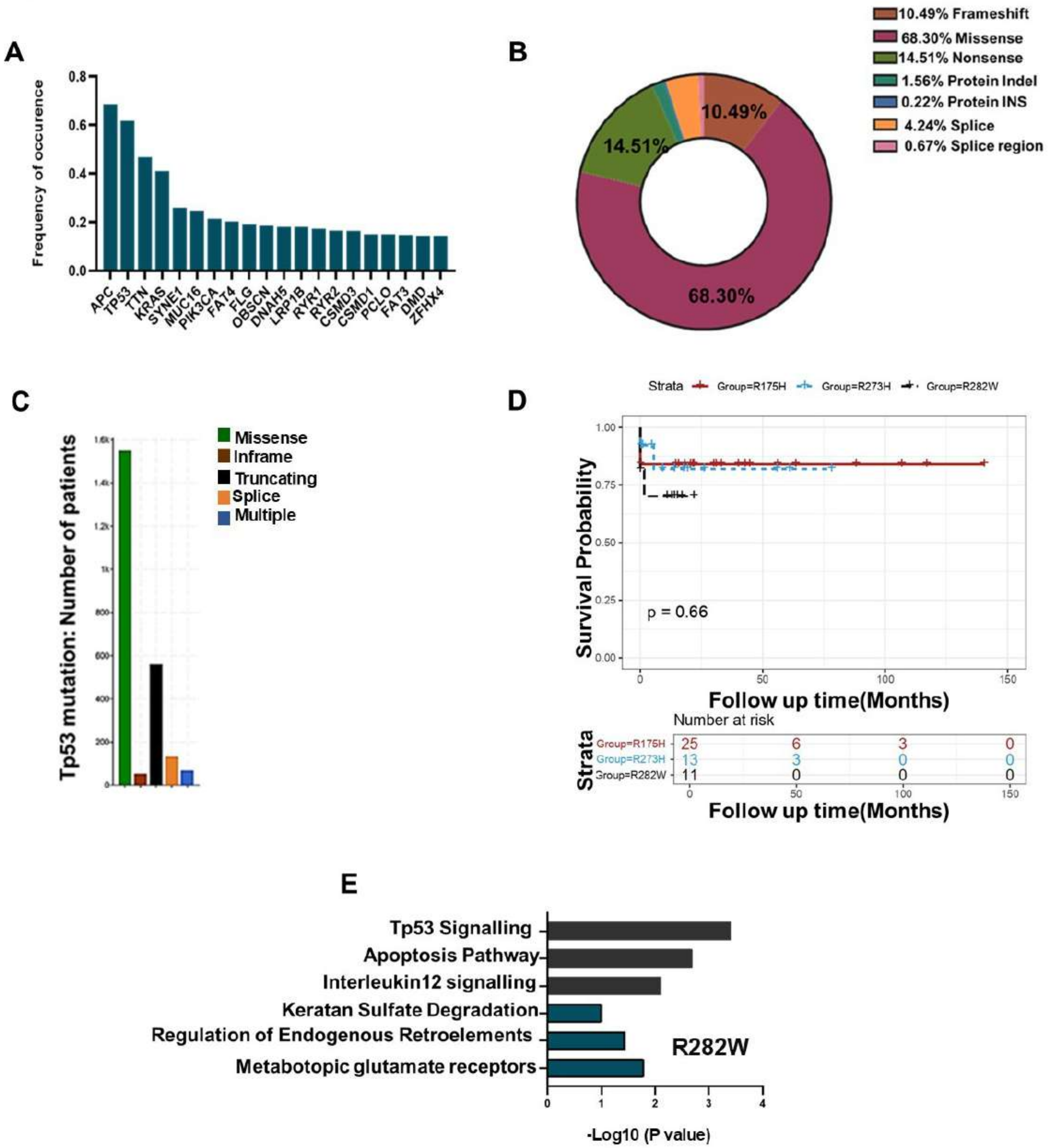
A
Frequency of occurence
0.8
0.6
0.4
0.2
0.0
APC
TP53
TTN
KRAS
SYNE1
MUC16
PIK3CA
FAT4
FLG
OBSCN
DNAH5
LRP1B
RYR1
RYR2
CSMD3
CSMD1
PCLO
FAT3
DMD
ZFHX4
B
10.49% Frameshift
68.30% Missense
14.51% Nonsense
1.56% Protein Indel
0.22% Protein INS
4.24% Splice
0.67% Splice region
10.49%
14.51%
68.30%
C
Tp53 mutation: Number of patients
Missense
Inframe
Truncating
Splice
Multiple
D
Strata
Group=R175H
Group=R273H
Group=R282W
Survival Probability
1.00
0.75
0.50
0.25
0.00
p = 0.66
0
50
100
150
Follow up time(Months)
Number at risk
Strata
Group=R175H 25 6 3 0
Group=R273H 13 3 0 0
Group=R282W 11 0 0 0
Follow up time(Months)
E
Tp53 Signalling
Apoptosis Pathway
Interleukin12 signalling
Keratan Sulfate Degradation
Regulation of Endogenous Retroelements
Metabotopic glutamate receptors
R282W
0
1
2
3
4
-Log10 (P value)

Fig S2

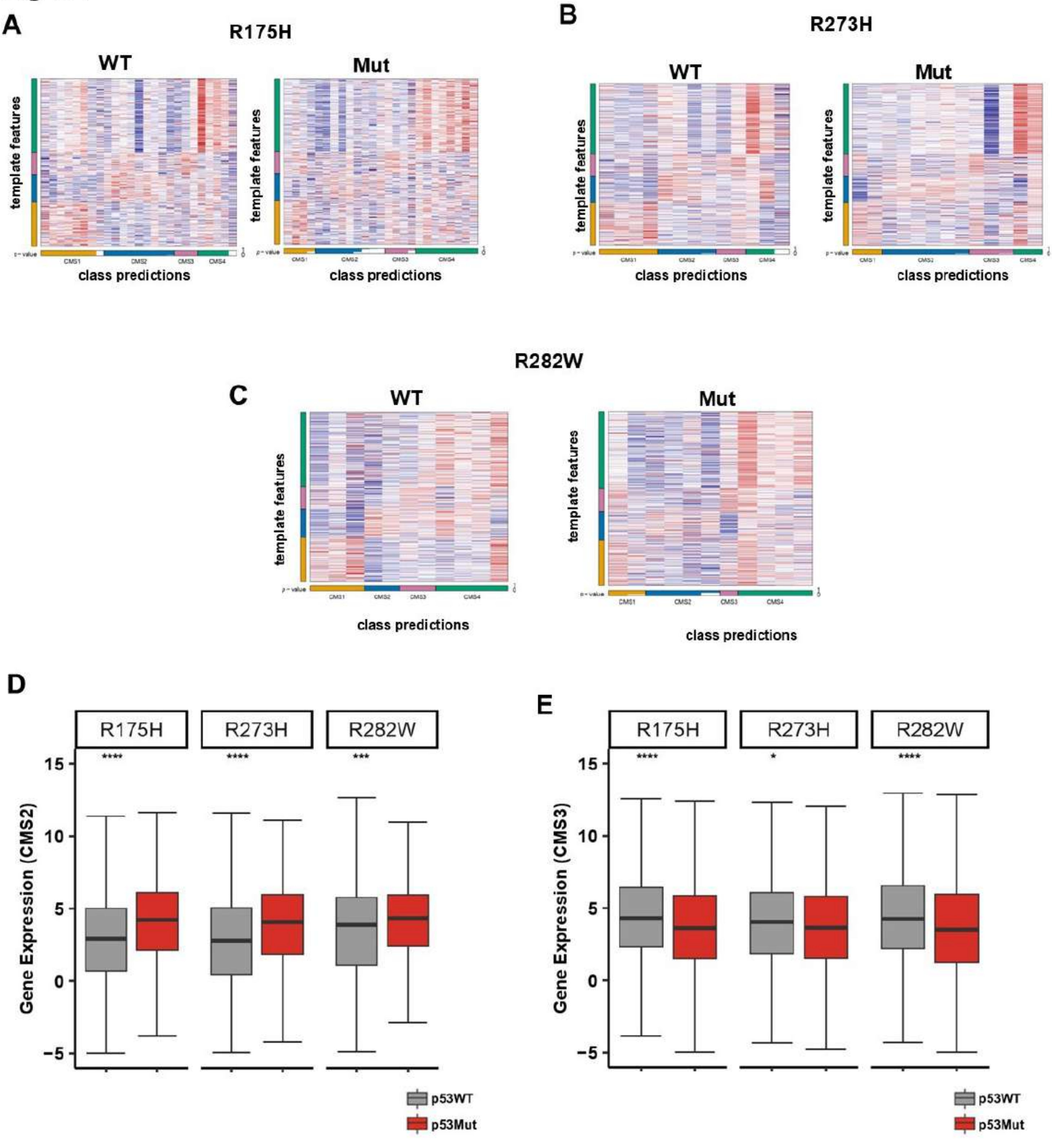

A
R175H
WT
Mut
template features
class predictions
CMS1
CMS2
CMS3
CMS4
B
R273H
C
R282W
D
E
R175H
R273H
R282W
****
***
*
Gene Expression (CMS2)
Gene Expression (CMS3)
15
10
5
0
-5
p53WT
p53Mut

Fig S3

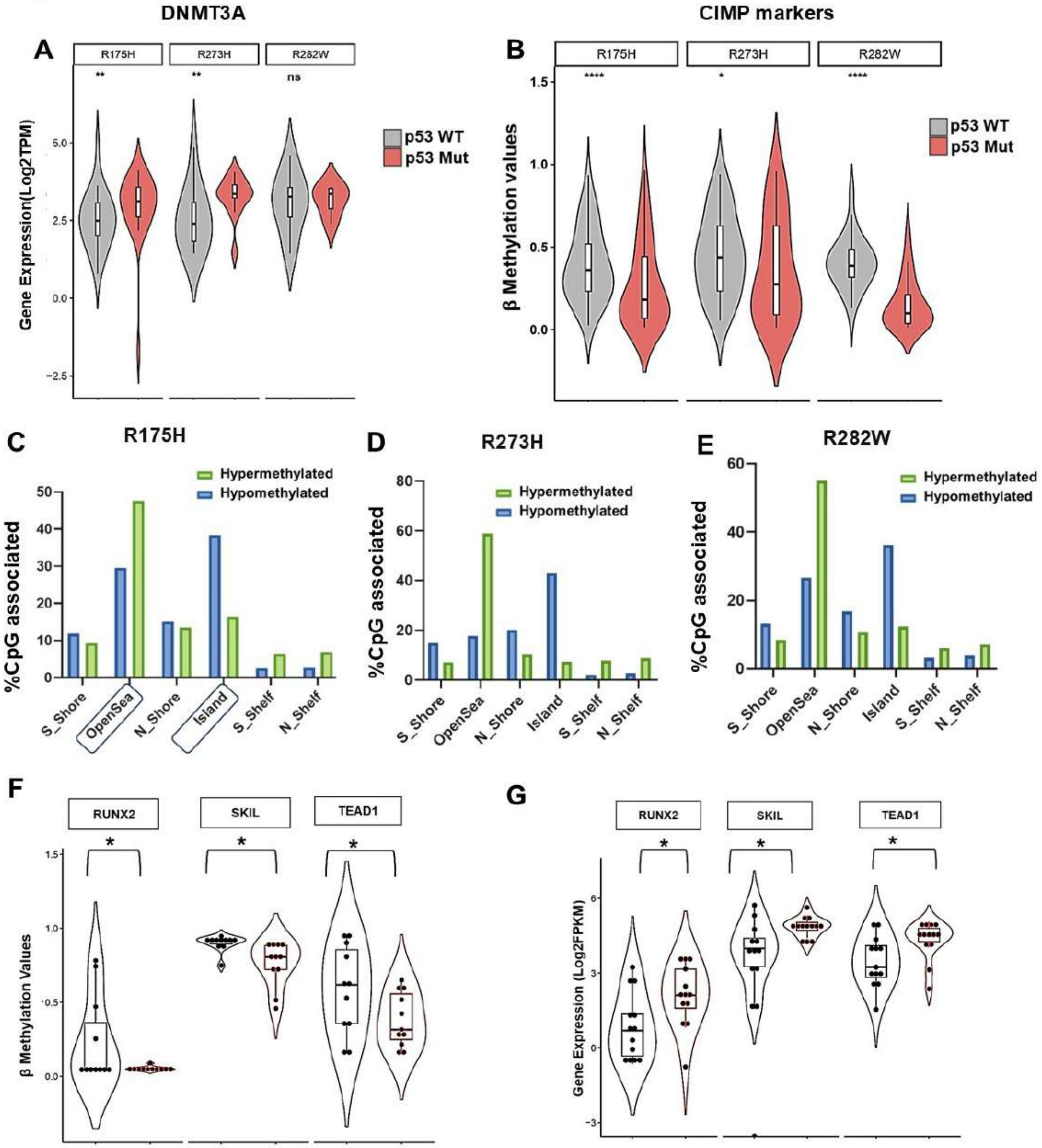

DNMT3A
CIMP markers
A
B
R175H
R273H
R282W
**
**
ns
****
*
****
Gene Expression(Log2TPM)
β Methylation values
p53 WT
p53 Mut
C
R175H
D
R273H
E
R282W
Hypermethylated
Hypomethylated
%CpG associated
S_Shore
OpenSea
N_Shore
Island
S_Shelf
N_Shelf
F
G
RUNX2
SKIL
TEAD1
β Methylation Values
Gene Expression (Log2FPKM)

### Fig S4

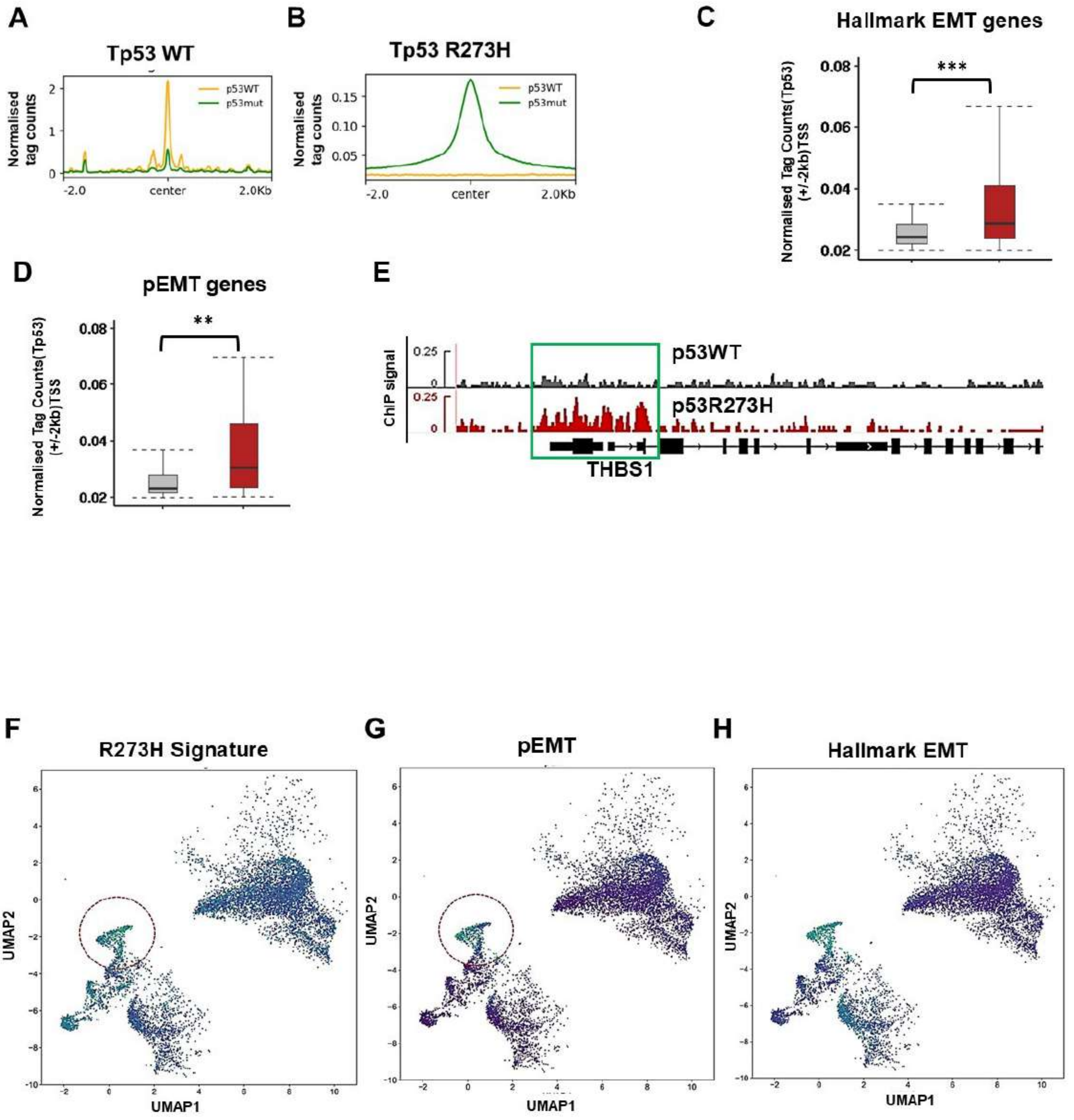

A
Tp53 WT
Normalised tag counts
p53WT
p53mut
-2.0
center
2.0Kb
B
Tp53 R273H
Normalised tag counts
p53WT
p53mut
0.15
0.10
0.05
-2.0
center
2.0Kb
C
Hallmark EMT genes
***
Normalised Tag Counts(Tp53) (+/-2kb)TSS
0.08
0.06
0.04
0.02
D
pEMT genes
**
Normalised Tag Counts(Tp53) (+/-2kb)TSS
0.08
0.06
0.04
0.02
E
ChIP signal
0.25
p53WT
p53R273H
THBS1
F
R273H Signature
G
pEMT
H
Hallmark EMT
UMAP1
UMAP2

### Supplementary Figure Legends

#### Fig S1: Stratification of colon tumours based on p53 mutational status

A. Barplot representing the status of Top 20 mutated genes in colon tumours. Tp53 is the second most mutated gene after APC.
B. Pie Chart representing prevalence of different Tp53 mutation in colon tumours as observed through TCGA-COAD. Missense mutations are the most prevalent mutations in Tp53.
C. Barplot representing the frequency of occurrence of different types of mutation in Tp53. Missense mutation is the most prevalent mutation followed by truncating mutation
D. Survival probability of three different p53 mutation in colon cancer. Kaplan Meier (KM) plot representing the survival profiles of top 3 prevalent Tp53 missense mutations. R273H tumours shows lower survivability as compared to R282W and R175H tumours. The X axis represents the survival time in months while the Y axis represents the survival probability

E. Column chart representing Gene Set Enrichment Analysis (GSEA) of the differentially expressed genes in p53R282W tumours (blue) and p53 WT (grey) (through reactome). R282W tumours show positive enrichment of metabotopic glutamate receptors.

#### Fig S2: Different p53 missense mutations show varied proportion of CMS subtypes based on associated gene expression

A. The consensus molecular subtypes (CMS) classification of CRC samples using CMScaller in p53 WT (left panel) and R175H (right panel). p53R175H associates with higher CMS4 proportion
B. The consensus molecular subtypes (CMS) classification of CRC samples using CMScaller in p53 WT (left panel) and R273H (right panel). p53R273H associates with higher CMS2 proportion
C. The consensus molecular subtypes (CMS) classification of CRC samples using CMScaller in p53 WT (left panel) and R282W (right panel). p53R282W associates with decrease in CMS1 proportion
D. Box plot representing the distribution of CMS2 gene expression in p53WT (grey) and p53mut(red). All three different p53 mutant tumours associate with significant increase in expression of CMS2 genes
E. Box plot representing the distribution of CMS2 gene expression in p53WT (grey) and p53mut(red). All three different p53 mutant tumours associate with significant decrease in expression of CMS3 genes

#### FigS3: Distinct methylation profiles around diverse genomic regions of CRC patients with p53 mutation

A. Box and violin plots representing the expression profile of the DNA methyl transferase DNMT3A in p53 WT (grey) and p53 mutant(red) tumours. R175H and R273H show significant higher expression of DNMT3A but not R282W tumors as compared to p53WT tumours.

B. Box and violin plot representing methylation values of CIMP markers in p53 WT (grey) and p53 mutant(red) tumours. All p53 mutant tumours associate with lower methylation values as compared to p53 WT tumours.
C. Column bar chart representing distribution of hypermethylated(green) and hypomethylated (blue) genomic regions with R175H mutation. Open sea regions correspond to hypermethylation profile, while CpG islands correspond to hypomethylation.
D. Column bar chart representing distribution of hypermethylated(green) and hypomethylated (blue) genomic regions with R273H mutation. Open sea regions correspond to hypermethylation profile, while CpG islands correspond to hypomethylation.
E. Column bar chart representing distribution of hypermethylated(green) and hypomethylated (blue) genomic regions with R273H mutation. Open sea regions correspond to hypermethylation profile, while CpG islands correspond to hypomethylation.

F. Box and violin plot representing the methylation levels of probes associated with RUNX2, SKIL and TEAD1 in p53WT tumours (black) and p53R273H tumours (red). R273H tumours associates with lower levels of methylation of probes associated with RUNX2, SKIL and TEAD1 as compared to p53 WT tumours.

G. Box and violin plot representing the expression profile of genes involved in YAP/TAZ signalling including RUNX2 and TEAD1 in p53WT tumours (black) and p53R273H tumours (red). R273H tumours shows significant upregulation of RUNX2, SKIL and TEAD1 as compared to p53WT tumours.

**Fig S4: Mutant p53 R273H enriches at partial/mesenchymal EMT states in colorectal cancer**

A. PlotProfile representing p53WT signal (yellow) and Tp53 R273H signal (green) at WT enriched regions (centered at +/-2kb peaks). WTp53 enriched regions correlates with higher WTp53 signal
B. PlotProfile representing p53WT signal (yellow) and Tp53 R273H signal (green) at R273H enriched regions (centered at +/-2kb peaks). R273H enriched regions correlates with higher R273H signal
C. Box plot representing normalized tag counts of WTp53(grey) and R273H(red) at hallmark EMT genes (+/2kb TSS). R273H shows significantly higher enrichment at the promoters of hallmark EMT gene sets.
D. Box plot representing normalized tag counts of WTp53(grey) and R273H(red) at pEMT genes (+/2kb TSS). R273H shows significantly higher enrichment at the promoters of pEMT gene sets.
E. IGV screenshot representing higher enrichment of p53R273H(red) at the promoters of THBS1(pEMT gene) as compared to p53WT(grey).

F-H.Signature enrichment visualized across tumour clusters: UMAP projections coloured by per-cell enrichment scores for (F) R273H transcriptional signature, (G) partial EMT (pEMT), and (H) Hallmark EMT. Colour scale (viridis): green = high enrichment, blue = low enrichment. The red dotted circle highlight a representative cluster (e.g., Tu1) where R273H enrichment co-occurs with elevated partial and full EMT programs, indicating mutant p53-associated phenotypic reprogramming.
